# An old inversion polymorphism involving giant mobile elements in an invasive fungal pathogen

**DOI:** 10.1101/2024.03.29.587348

**Authors:** Fanny E. Hartmann, Ricardo C. Rodriguez de la Vega, Arthur Demené, Thomas Badet, Jean-Philippe Vernadet, Quentin Rougemont, Amandine Labat, Alodie Snirc, Lea Stauber, Daniel Croll, Simone Prospero, Cyril Dutech, Tatiana Giraud

## Abstract

Recombination suppression can evolve in sex or mating-type chromosomes, or in autosomal supergenes, with different haplotypes being maintained by balancing selection. In the invasive chestnut blight fungus *Cryphonectria parasitica*, a genomic region was suggested to lack recombination and to be partially linked to the mating-type (MAT) locus based on segregation analyses. Using hundreds of available *C. parasitica* genomes and generating new high-quality genome assemblies, we show that a ca. 1.2 Mb genomic region proximal to the mating-type locus lacks recombination, with the segregation of two highly differentiated haplotypes in balanced proportions in invasive populations. High-quality genome assemblies further revealed an inversion in one of the haplotypes in the invaded range. The two haplotypes were estimated to have diverged 1.5 million years ago, and each harboured specific genes, some of which likely belonging to *Starship* elements, that are large mobile elements, mobilized by tyrosine recombinases, able to move accessory genes, and involved in adaptation in multiple fungi. The MAT-proximal region carried genes upregulated under virus infection or vegetative incompatibility reaction. In the native range, the MAT-proximal region also appeared to have a different evolutionary history than the rest of the genome. In all continents, the MAT-Proximal region was enriched in non-synonymous substitutions, in gene presence/absence polymorphism, in tyrosine recombinases and in transposable elements. This study thus sheds light on a case of a large non-recombining region partially linked to a mating compatibility locus, with likely balancing selection maintaining differentiated haplotypes, possibly involved in adaptation in a devastating tree pathogen.

## Introduction

Recombination increases the efficiency of selection, the purging of deleterious alleles and the generation of potentially beneficial allelic combinations (Otto and Lenormand 2002). However, recombination can also break up beneficial allelic combinations and, therefore, can be selected against, generating supergenes, i.e., large regions without recombination encompassing multiple genes (Schwander et al. 2014). This is often the case in genomic regions controlling mating compatibility, such as regions determining sex, mating type or self-incompatibility in plants, animals, fungi, algae and oomycetes (Charlesworth et al. 2005; Bergero and Charlesworth 2009; Umen 2011; Charlesworth 2016; Dussert et al. 2020; Hartmann et al. 2021). Such recombination suppression keeps alleles linked across different genes that prevent self-compatibility or intermediate sexual phenotypes, and can sometimes extend away from sex-determining genes or mating-type loci (Bergero and Charlesworth 2009; Furman et al. 2020; Hartmann et al. 2021). Even beyond sex and mating-type chromosomes, there is an increasing number of supergene reports in autosomes (Schwander et al. 2014) with highly differentiated haplotypes maintained by balancing selection. Striking examples include supergenes controlling social structure in ants, wing color patterns in butterflies, reproductive morphs in birds or host-parasite interactions (Küpper et al. 2016; Yan et al. 2020; Jay et al. 2021; Fredericksen et al. 2023).

In the long term, however, because of less efficient selection, genomic regions without recombination show signs of degeneration, such as transposable element accumulation, rearrangements, non-synonymous substitutions, decreased gene expression, reduced frequency of optimal codons and gene losses (Bachtrog 2013; Carpentier et al. 2022; Duhamel et al. 2023). The non-recombining regions can nevertheless be maintained by balancing selection, with different allelic combinations having contrasting advantages in different situations or experiencing negative frequency dependent selection (Jay et al. 2021; Berdan et al. 2022). The evolutionary mechanisms leading to recombination suppression and allowing their persistence remain however poorly known (Jay, D. Jeffries, Hartmann, et al. 2024). Reporting more diverse constellations of recombination suppression in a variety of organisms with contrasting life-history traits is important for understanding the general patterns of recombination suppression, their evolution and maintenance (Ironside 2010; Charlesworth 2016; Furman et al. 2020; Hartmann et al. 2021).

In the invasive chestnut blight fungal pathogen *Cryphonectria parasitica*, mating compatibility is determined by a mating-type locus that displays two alleles, called *MAT-1* and *MAT-2,* that are both permanently heterozygous in the diploid and dikaryotic stages (McGuire et al. 2001). A large region without recombination was suggested to occur near the mating-type locus, although not completely linked to it (*i.e.,* at 3.9 cM of the mating-type locus), based on segregation of RAPD and RFLP markers in three crosses involving five different parents, from the USA, Japan or Italy (Kubisiak and Milgroom 2006a). Recent genome-wide association studies in a worldwide collection and in local *C. parasitica* populations in southern Switzerland found significant association of SNPs with the mating-type locus across a large region (>1 Mb), further suggesting the existence of reduced recombination in this region (Stauber et al. 2021; Stauber et al. 2022). This genomic region was enriched in SNPs, transposable elements and copy-number variants, such as deletions, which are consistent with sequence degeneration and a lack of recombination (Stauber et al. 2021), and which prevented its assembly so far.

The evolutionary history of the species is well documented as it is an invasive and highly damaging pathogen, having almost caused the extinction of American chestnut (*Castanea dentata*) in North America (Anagnostakis 1987). From its center of origin in Asia and its original hosts, the <u>Chinese chestnut</u> (*Ca. mollissima*) and the Japanese chestnut (*Ca. crenata*), *C. parasitica* has first invaded North America, killing millions of American chestnuts (*Ca. dentata*). In Europe, at least two distinct introduction events occurred, on the <u>European chestnut</u>(*Ca. sativa*): one introduction from North America to Italy, and one directly from Asia, probably to the Pyrenees Mountains. European strains will be therefore hereafter referred to as invasive European strains introduced from North America or invasive European strains introduced directly from Asia. The chestnut blight symptoms have been less severe in Europe than in North America, due to lower susceptibility of *Ca. sativa* and a virus infecting *C. parasitica* and causing hypovirulence (Dutech et al. 2010; Dutech et al. 2012). The genetic determinants of the adaptation of the pathogen to its new environments and hosts remain largely unknown (Lovat and Donnelly 2019).

Lineages with an apparently clonal structure have been identified in invasive populations while sexually reproducing populations occur both in the native and invaded ranges (Dutech et al. 2012; Stauber et al. 2021). The lineages with a predominant clonal structure likely still undergo rare sex events, as shown by the presence of the two mating-type alleles in most of them (Demené et al. 2019). Furthermore, although this ascomycete fungus is mostly found as haploid mycelia, some isolates have been reported to be heterokaryotic at the mating-type locus in several invasive European and North American populations, i.e., with cells carrying different nuclei, of the opposite mating types *MAT-1* and *MAT-2* (McGuire et al. 2004; McGuire et al. 2005; Dutech et al. 2010; Stauber et al. 2021; Stauber et al. 2022).

Here, we therefore studied the occurrence of recombination suppression in the genomic region proximal to the mating-type locus in *C. parasitica,* using the previously published genomes and further generated six new high-quality long-read based genome assemblies of strains from the native and invaded range of the pathogen. More specifically, we tested whether recombination was completely suppressed in the genomic region proximal to the mating-type locus in *C. parasitica*. We first analyze sexually reproducing invasive European populations, for which extensive genomic datasets are available in local populations. We assessed, in these populations, the level of differentiation between haplotypes, as well as the frequency of the two non-recombining haplotypes and their association with mating types. We investigated whether other regions of the genome showed reduced recombination rates and whether genomic footprints of degeneration were present in the non-recombining region, such as genomic rearrangements, non-synonymous substitutions and transposable element accumulation. We also tested whether this non-recombining region has been gradually expanding, estimated the age of recombination suppression and investigated whether the predicted gene functions in the MAT-proximal region could help understanding the evolutionary cause for recombination suppression, polymorphism maintenance and partial linkage to the mating-type locus. We looked in particular for *Starships*, that are giant mobile elements recently discovered in ascomycete fungi and able to move accessory genes as cargo within and between genomes (Gluck-Thaler et al. 2022; Urquhart et al. 2024). These cargo genes can be involved in adaptation (Gluck-Thaler et al. 2022; Urquhart et al. 2024), as shown in fungal pathogens of coffee (Peck et al. 2023) and of wheat (Bucknell et al. 2024; Tralamazza et al. 2024), as well as in molds used for making cheeses (Cheeseman et al. 2014; Ropars et al. 2015) and those maturing dry-cured meat (Lo et al. 2023). *Starships* are characterized by their captain, a tyrosine recombinase with a DUF3435 domain, being the first gene at the 5’ edge of the elements and allowing their excision and insertion. *Starships* can contain “cargo” genes, highly variable in nature and number among *Starship* elements (Urquhart et al. 2024). *Starship* diversity is partitioned into 11 major families, targeting specific genomic niches for their insertions, e.g., AT-rich regions (Gluck-Thaler et Vogan, 2024). We then extended the analyses to other populations with less extensive genomic resources, i.e. other invasive populations and native Asian populations, investigating the presence of recombination suppression footprints and the occurrence of the two differentiated haplotypes worldwide, the presence of *Starships*, as well as the possibility of introgression, using available genomes of closely related species, pathogenic and non-pathogenic (Stauber et al. 2021). Finally, we used available expression data to investigate whether the MAT-Proximal region carried genes upregulated under infection by the virus responsible for hypovirulence or during vegetative incompatibility reactions, a phenomenon avoiding hyphal fusions between individuals, considered to protect against virus transmission.

## Results

### Footprints of recombination suppression in a large region (> 1 Mb) proximal to the mating-type locus in sexually reproducing invasive populations

We first performed population genomic analyses on available genome sequences within local populations sampled in southern Switzerland over two temporal frames (early 1990 and 2019), and within the more broadly geographically distributed CL1 cluster in central and southeastern Europe, both having a recombining genetic structure and being invasive European populations introduced from North America (Table S1 (Stauber et al. 2021)). We studied only monokaryotic strains, i.e. having either *MAT-1* or *MAT-2* in their genomes, for phasing haplotypes and performed stringent SNPs filtering by masking repeats on the EP155 reference genome (Crouch et al. 2020) and removing missing data and rare variants to ensure robustness of our analyses. We identified 8,900 SNPs segregating among 71 strains of the 1990 Swiss population (35 MAT-1; 36 MAT-2; Table S1), 9,646 SNPs segregating among 62 strains of the Swiss 2019 population (20 MAT-1; 42 MAT-2; Table S1) and 15,104 SNPs segregating among 88 strains of the CL1 cluster (41 MAT-1; 47 MAT-2; Table S1), which represent an average density of about 20 SNPs/100 kb, in agreement with previous studies in this invasive fungus (Stauber et al. 2021; Stauber et al. 2022). We investigated the genome-wide linkage disequilibrium (LD) landscape and looked for large blocks (>1 Mb) of high LD (r2>0.9) among SNPs within contigs, to look for signatures of reduced recombination along the genome. LD indeed decays by half across a few 100 kb on average in *C. parasitica* in recombining regions of the genome (Demené et al. 2019). Therefore, high LD beyond >1 Mb represents strong evidence of recombination cessation and the SNP density is sufficient to study such LD variation.

We found a large block of high LD on the chromosome called scaffold_2 of the EP155 genome in the three populations, corresponding to the mating-type chromosome (see Fig 1 for the CL1 population, the pattern in the other populations being similar). The high-LD block was located near the mating-type locus but did not encompass the mating-type locus itself and will be called hereafter the “MAT-proximal region”. In the 1990 and 2019 Swiss populations, SNPs between 0.493 Mb and 1.720 Mb in the mating-type chromosome were all in high LD (with r^2^>0.9 between SNPs distant by up to 1 Mb; in this region, mean r^2^ of all SNP pairs in the Swiss 1990 population = 0.89; mean r^2^ of all SNP pairs in the Swiss 2019 population = 0.83; Fig S1). In the CL1 cluster, the large high-LD block was also present, but was split into two blocks, a large one and a smaller, peripheral one (orange arrows in Fig 1). We indeed detected values of r^2^>0.9 between SNPs distant by up to 1 Mb, between 0.532 Mb and 1.535 Mb (mean r^2^ of all SNP pairs = 0.94). Between 1.535 and 1.717 Mb, SNPs were also in high LD (mean r^2^ = 0.84). The LD between the two high-LD blocks was lower (r^2^= 0.59) than within blocks but was still higher than elsewhere along the genome (orange rectangle at the right border of the red triangle; Fig 1). A similar pattern also existed in the Swiss populations (orange rectangle at the right border of the red triangle; Fig S1), although less marked. This is likely due to rare recombination events at one specific locus near the edge of the large fully non-recombining region, lowering the LD level in populations between the two parts of the otherwise fully non-recombining region. The mating-type locus was located at 1.737 Mb in the EP155 genome, distant by 17 kb and 20 kb from the LD block in the Swiss populations and CL1 cluster, respectively. SNP density was on average 48 SNPs/100 kb in the high-LD block region, i.e. higher than in the rest of the chromosome (Fig S2C). When sampling SNPs distant of at least 50 kb in this region to have the same SNP density all over the chromosome, we also detected the high-LD block, which indicates that the SNP density did not generated biases in the LD pattern (Fig S2D).

**Figure 1:**
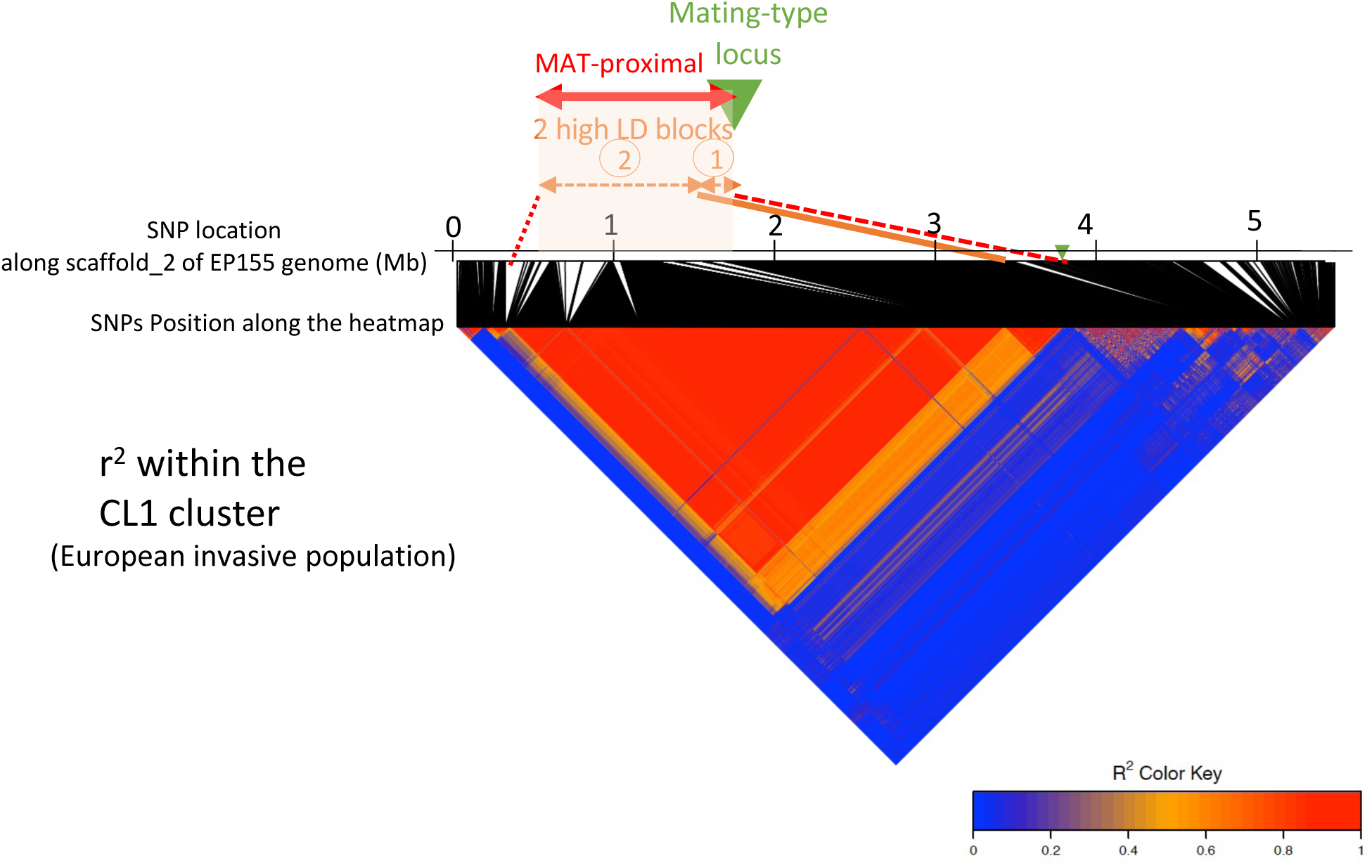
Linkage disequilibrium (LD) analysis along the contig carrying the mating-type locus in a *Cryphonectria parasitica* European invasive population. LD heatmaps using single nucleotide polymorphisms (SNPs; n=3815) located on the contig carrying the mating-type locus (scaffold_2 of the EP155 genome) in the CL1 genetic cluster (European invasive population introduced from North America); pairs of SNPs with high LD, i.e. r^2^ >0.9, correspond to the red triangle. The mating-type locus location is shown with a green triangle and the MAT-proximal region lacking recombination is shown with a red arrow. The two high-LD blocks within the MAT-proximal region are shown with orange arrows. SNPs at the limit of the MAT-proximal region and the two high LD blocks were manually highlighted with red dotted lines and an orange line.

We found no other regions of the genomes with r^2^>0.9 between SNPs across such a large genomic distance in any population. On scaffold_6 in the CL1 cluster, two smaller SNP blocks had a r^2^>0.9 between each other despite being distant (Fig S2C; between SNPs blocks at 3-20 kb and 2.151-2.278 kb), which is likely due to the major intra-scaffold translocation described in this region between the EP155 and ESM15 strains (Demené et al. 2022). The maximum size of the blocks with r²>0.9 on other scaffolds ranged from 135 to 540 kb in the CL1 cluster, as previously described (Demené et al. 2019). The average distance between SNPs in high LD (r²>0.9) across the genome was higher on the scaffold_2 than on other scaffolds (in the CL1 cluster, pairwise Wilcoxon test p-value < 2e-16 with Bonferroni correction). Furthermore, the proportion of SNPs in high LD (r^2^>0.9) across the genome was also much higher on the scaffold_2 than on other scaffolds due to the high LD in the MAT-proximal region (Fig S2D, Table S2). Therefore, the large (1Mb) and localized block with maximal LD values proximal to the mating-type locus both in the Swiss populations and the CL1 genetic cluster stands out as exceptional in the genome and indicates full recombination cessation, which supports previous inferences from progeny segregation analyses (Kubisiak and Milgroom 2006b). Indeed, even low rates of recombination homogenize allelic frequencies and prevents LD building (Dufresnes et al. 2015). The MAT-proximal non-recombining region was actually larger (1.23 Mb), but with likely rare recombination events at one precise locus near its edge, lowering LD between its two fully non-recombining parts.

Consistent with a lack of recombination, the MAT-proximal region formed two genetically highly differentiated haplotypes in these invasive Swiss and CL1 populations (Fig 2A; Fig S3 A(1)-C(1)), as shown by the two clusters on the principal component analysis (PCA) using only SNPs located in the MAT-proximal region, while no structure was detected in the rest of the genome (Fig 2B-C; Fig S3A(2)-C(2)). The neighbor-net networks further supported the existence of two differentiated haplotypes in the MAT-Proximal region, contrasting with an otherwise recombining structure genome-wide (Fig 2D-E; Fig S3B-D). In the Swiss 2019 population, a few reticulations between haplotypes were found and three strains (LU3, Nov10, Nov4) appeared to have an intermediate sequence in the MAT-Proximal region between the two haplotypes (Fig S3D). This intermediate haplotype was also found in a few other invasive strains introduced directly from Asia (see below).

**Figure 2:**
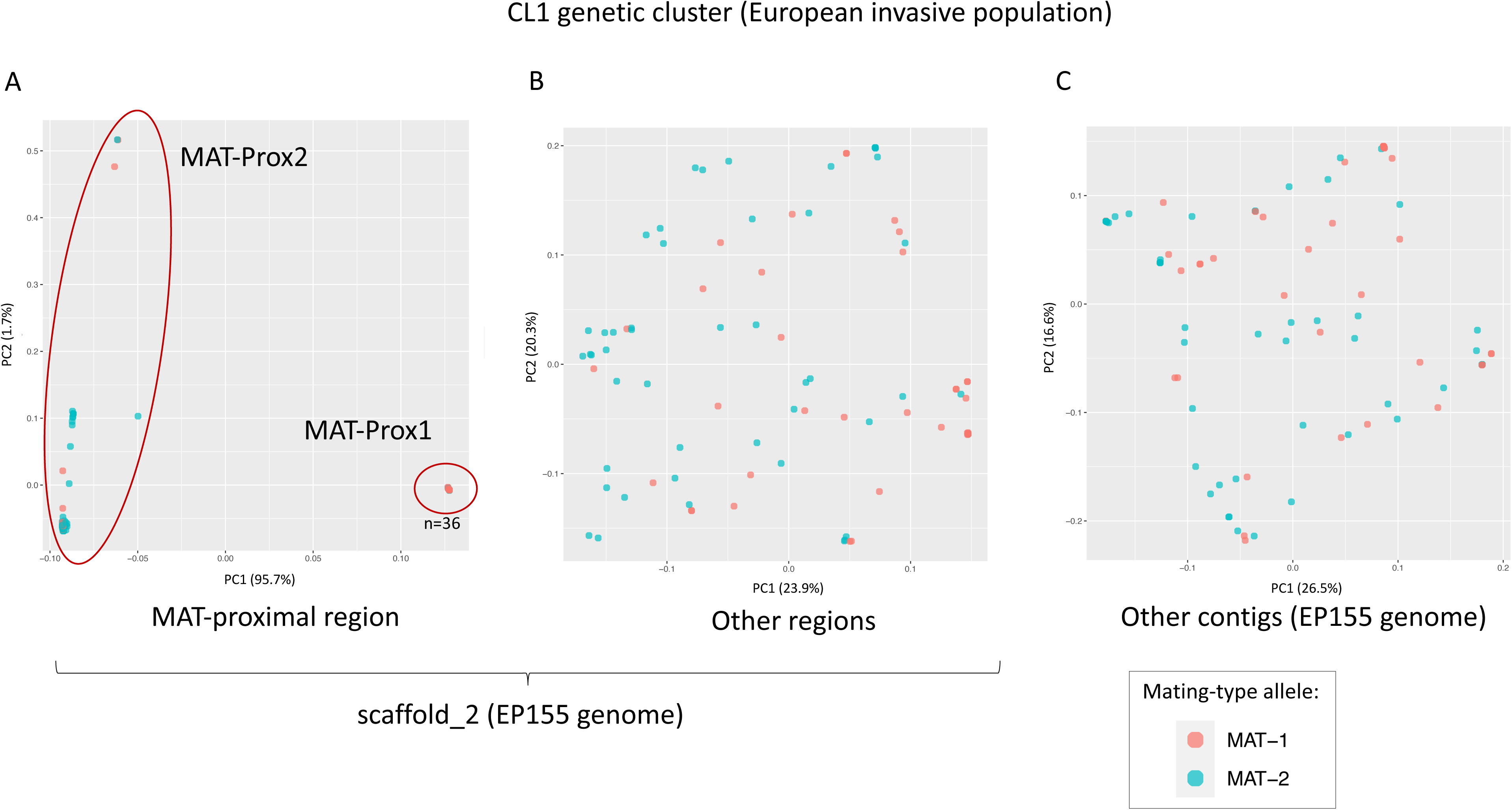
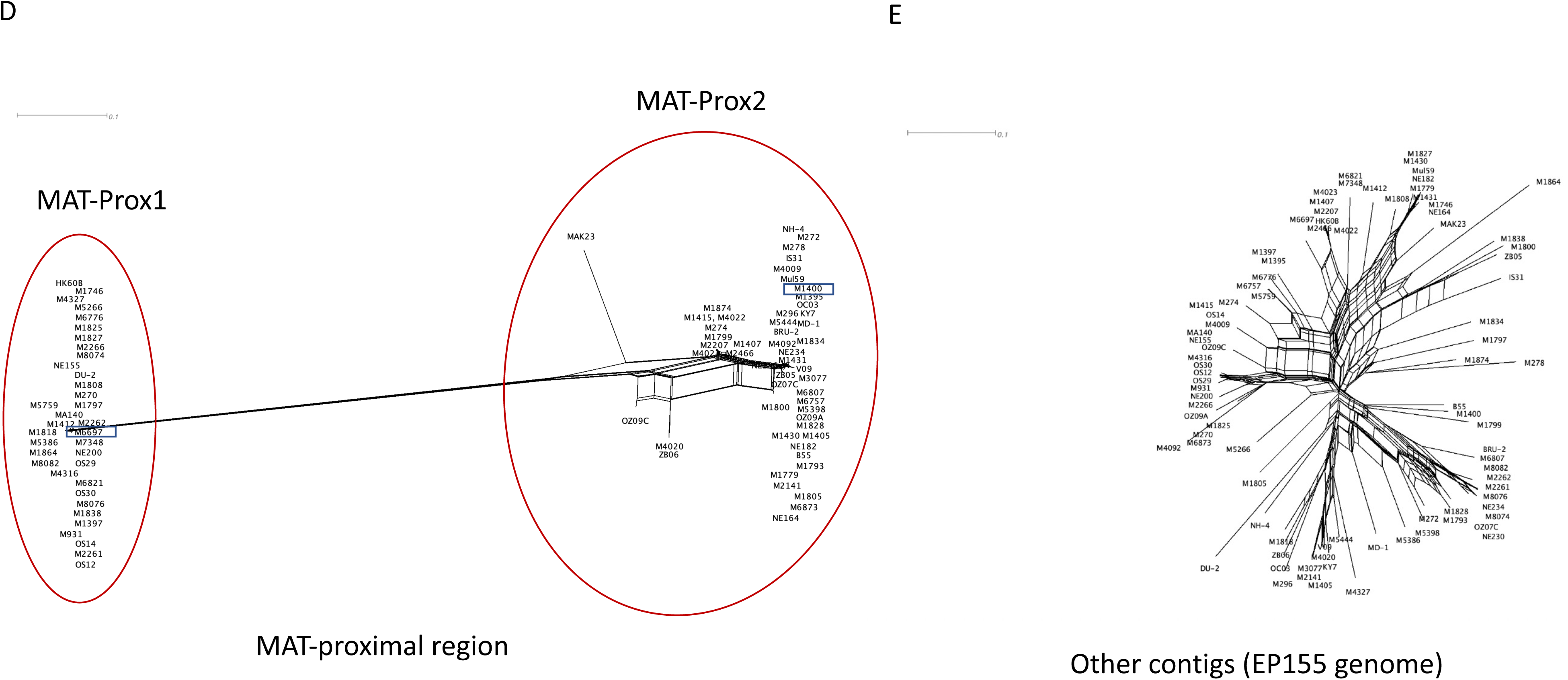
Genetic structure using single nucleotide polymorphisms (SNPs) in the MAT-proximal region lacking recombination and other genomic regions in a *Cryphonectria parasitica* European invasive population. A-C. Principal component analysis (PCA). Two principal components are presented. Percentage of variance explained by each PC is indicated into brackets. Strains are colored according to their mating type (*MAT-1* or *MAT-2*). **D-E.** Neighbor-net network from a SplitsTree analysis. These analyses were performed in the CL1 genetic cluster (European invasive populations introduced from North America) based on: **A and D.** SNPs (n=2,220) located within the MAT-proximal region along the contig carrying the mating-type locus (scaffold_2 of the EP155 genome).; **B.** SNPs (n=1,595) located in other regions along the contig carrying the mating-type locus; **C and E** SNPs (n=11,289) located on other contigs of the EP155 genome. On panels A and D, the two clusters corresponding to the MAT-Prox1 and MAT-Prox2 haplotypes are shown with red circles. The identified haplotype of each strain is indicated in Table S1. The number of strains within the MAT-Prox1 haplotype is indicated on panel A by the letter n. The newly sequenced M1400 and M6697 strains are highlighted with a black rectangle

The two genetic clusters in the MAT-proximal region appeared associated with mating types in the CL1 and Swiss 1990 recombining populations (Figs 2A and D; Fig S3, Table 1). Indeed, the distribution of mating types between the two PCA clusters strongly deviated from expectations under random association (Table 1; chi-squared test: χ² = 33.3; p-value = 5.876e-08 in CL1; χ² = 18.76; p-value = 8.441e-05 in Swiss 1990). We thus hereafter call the two differentiated haplotypes MAT-Prox1 and MAT-Prox2, referring to the mating-type association. However, the association between mating types and MAT-Proximal haplotypes was not complete, with for example only 83% of MAT1-1 strains carrying the MAT-Prox1 haplotype in CL1. The two MAT-Proximal haplotypes were not significantly associated with mating types in the Swiss 2019 recombining populations (Table 1; chi-squared test: χ² = 3.8427; p-value = 0.1464). The EP155 genome used as reference for SNP calling above carried the MAT-Prox1 haplotype but the MAT1-2 allele.

**Table 1:** Distribution of mating types (*MAT-1* and *MAT-2*) among non-recombining haplotypes of the MAT-proximal region in the European invasive *Cryphonectria parasitica* CL1 genetic cluster and the Swiss populations. MAT-Prox1 and MAT-Prox2 haplotypes were defined based on the clusters of the principal component analysis.

|  | Population | CL1 genetic cluster |  | Swiss 1990 population |  | Swiss 2019 population |  |
| --- | --- | --- | --- | --- | --- | --- | --- |
|  | Haplotype | MAT-Prox1 | MAT-Prox2 | MAT-Prox1 | MAT-Prox2 | MAT-Prox1 | MAT-Prox2 |
| Number of strains<br>in each PCA cluster | total | 36 | 52 | 20 | 51 | 8 | 54 |
|  | *MAT-1* | 30 | 11 | 18 | 17 | 5 | 15 |
|  | *MAT-2* | 6 | 41 | 2 | 34 | 3 | 39 |
| % of strains | *MAT-1* | 83 | 21 | 90 | 33 | 63 | 28 |
| in each PCA cluster | *MAT-2* | 17 | 79 | 10 | 67 | 38 | 72 |

### Inversion in the non-recombining MAT-proximal region

To investigate whether rearrangements between haplotypes were present in the MAT-proximal region, we sequenced the genome, using PacBio HiFi, of two strains with alternative MAT-proximal haplotypes: M1400 (MAT-2; MAT-Prox2) and M6697 (MAT-1; MAT-Prox1), originating from the Swiss population in Gnosca and belonging to the CL1 cluster, i.e. European invasive populations introduced from North America (Stauber et al. 2021; Stauber et al. 2022). We built high-quality genome assemblies: statistics of the assemblies were in the same range as those of the genome assemblies of the ESM15 and EP155 strains previously sequenced (Table S3; Demené et al 2022; Crouch et al 2020). The mating-type chromosome could be assembled as a single contig for the first time, in the M1400 genome, likely corresponding to a full chromosome, and was assembled into two contigs in the M6697 genome. We therefore used the M1400 genome as a reference for subsequent analyses. To identify the location of the MAT-proximal region in the M1400 genome, we computed LD by mapping the reads of the 1990 Swiss population to the new M1400 genome. We found the large block (> 1 Mb) of high LD (r^2^>0.9) between 7,285,137 and 8,828,934 bp (red arrow; Fig. S4), showing that calling SNPs on either a MAT-Prox1 or MAT-Prox2 haplotype yielded similar LD patterns. The MAT-proximal region was also located 20 kb away from the mating-type locus (located at 7.265 Mb) and was 1.540 Mb long using the M1400 genome as reference. Consistent with the results using the EP155 genome as reference, we found that the high-LD region was divided into two higher-LD blocks near the edge at 7.392 Mb (orange arrows in Fig. S4).

The two new high-quality genome assemblies from the invasive Swiss population (introduced from North America) revealed an inversion between the two haplotypes in the MAT-proximal region (Fig 3). The two newly sequenced PacBio genomes were indeed collinear except for the mating-type chromosome (Fig S5A-B-C), where we found a large region (>1 Mb) that seem inverted between the M1400 and M6697 genomes (Fig 3; Fig S5A-B-C). Breakpoints of the inversion were located at ca. 7.455 and 8.563 Mb of the tig00000001 contig in the M1400 genome and at ca. 3.380 and 4.185 Mb of the tig00000060 contig in the M16697 genome. The region affected by the inversion was ca. 300 kb smaller in the M6697 genome than in the M1400 genome, while the whole MAT-proximal region was ca. 100 kb smaller. The split into the two high-LD blocks in natural populations reported above was outside of the inversion, i.e. 70 kb from the inversion breakpoint (orange arrows in Fig S4). LD in natural populations was higher within the inversion (median value of r^2^ = 1) than in other regions of the MAT-proximal region (median value of r^2^ = 0.64; Wilcoxon rank sum test with continuity correction; W = 4.8714e+10; p-value < 2e-16).

**Figure 3:**
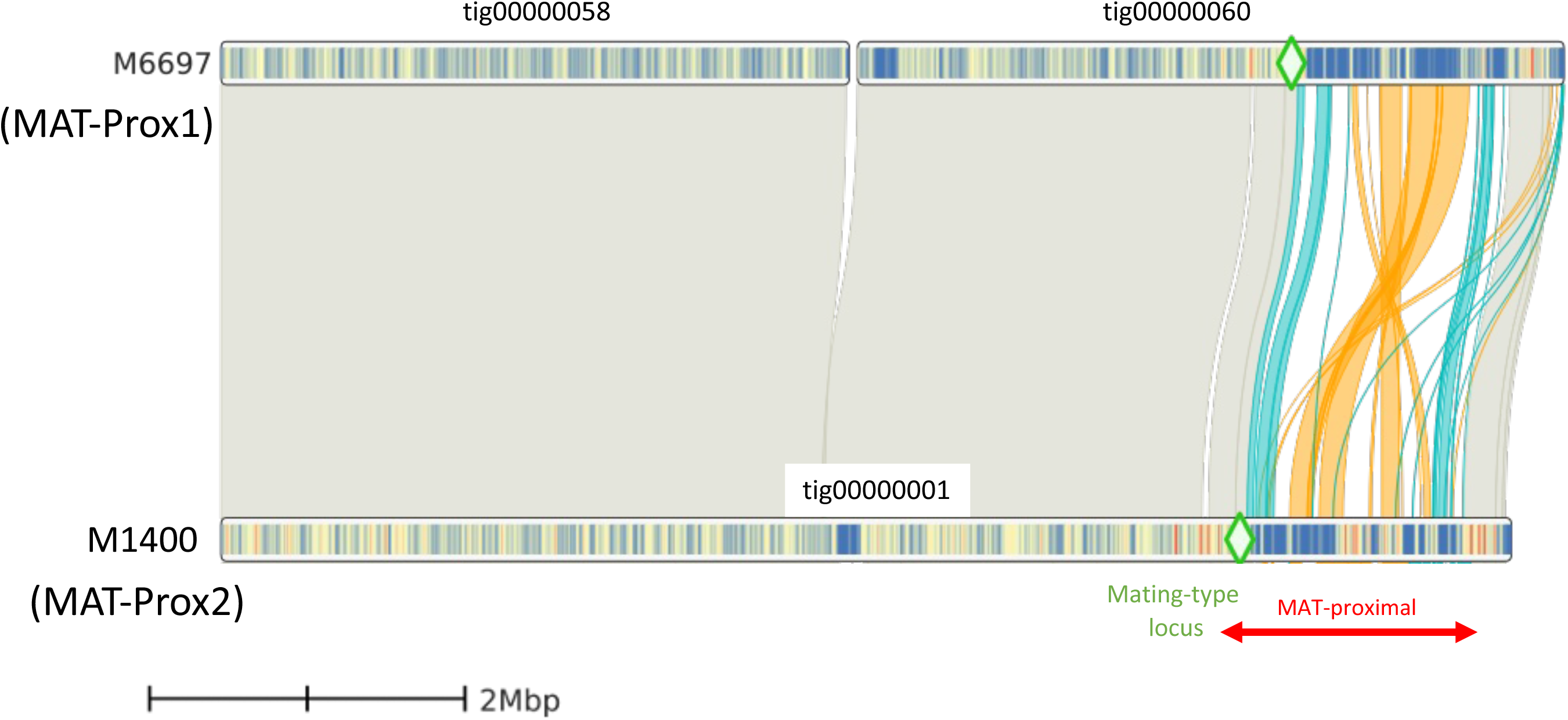
Synteny and rearrangements between the two newly sequenced genomes of European invasive strains introduced from North America (M1400 and M6697) in the chromosome carrying the MAT-proximal region lacking recombination in *Cryphonectria parasitica*. Blue links show colinear regions and orange links show inverted regions in the MAT-proximal region. Grey links show other regions. The mating-type locus is located with a green diamond. The MAT-proximal region defined from LD analyses is indicated with red arrows. Gene density tracks is shown with a color gradient (blue with low density, orange with high density).

We looked for centromeres, as the non-recombining regions near the mating-type locus in other ascomycetes, when they occur, either capture the centromere (Menkis et al. 2008; Sun et al. 2017), or are associated to the occurrence of a single crossing-over between the centromere and the mating-type locus (Grognet et al. 2014; Hartmann et al. 2021; Vittorelli et al. 2022). The centromere in *C. parasitica* may be at 4.380-4.536 Mb on the mating-type chromosome in the M1400 genome, as we detected here a peak in TE density and a drop in GC content (Fig 4A-B). A dotplot of repeats in this region also presented a pattern typical of centromeres (Fig S5D). The MAT-proximal region and the inversion thus did not include the putative centromere and was instead located about 2.9 Mb away of the putative centromere.

**Figure 4:**
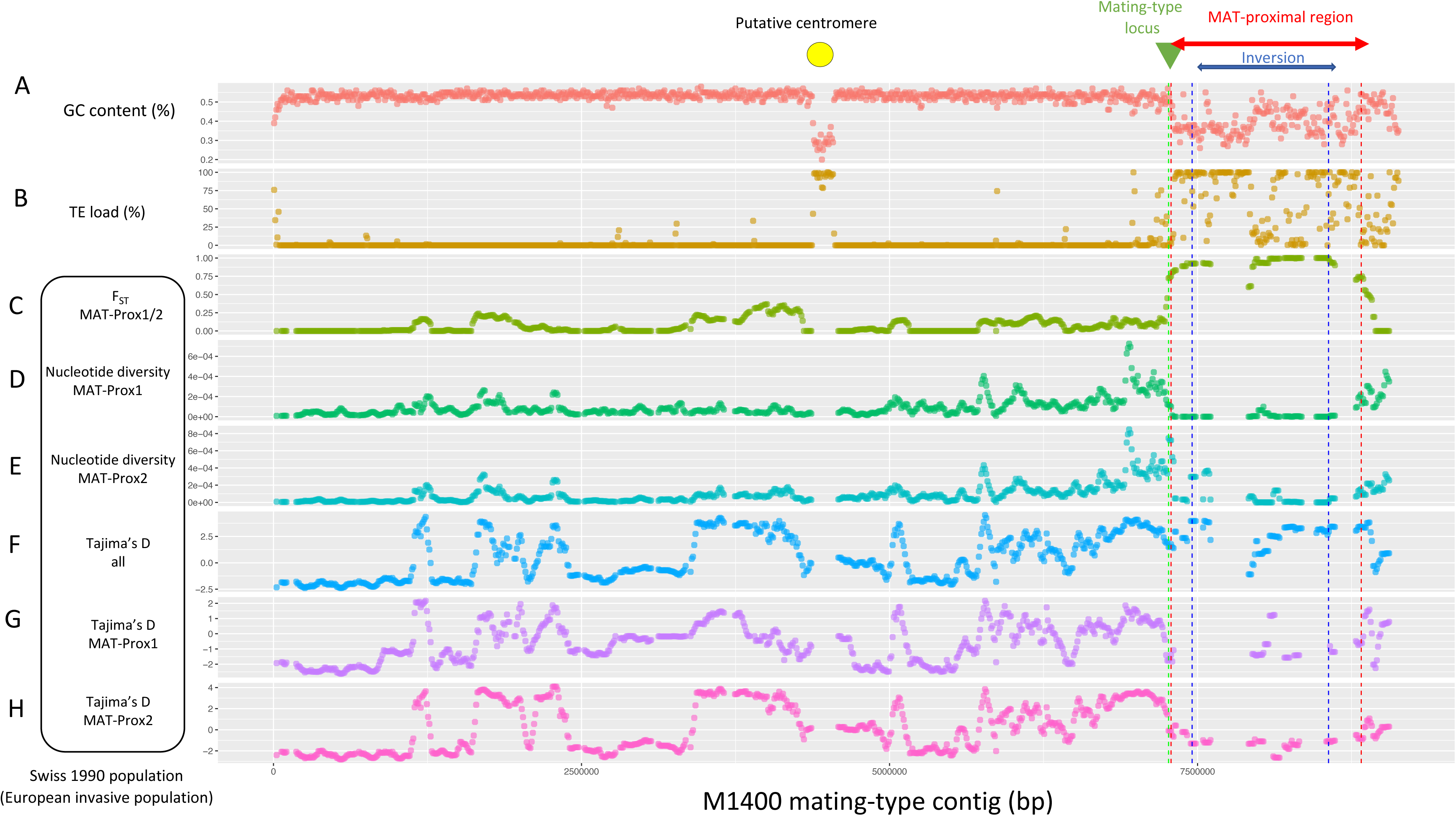
Genetic diversity and divergence between non-recombining haplotypes of the MAT-proximal region along the contig carrying the mating-type locus in a *Cryphonectria parasitica* European invasive population. A-B. GC content (A) and transposable element densities (B) along the contig carrying the mating-type locus (tig_0000001) of the M1400 *C. parasitica* genome. C. Relative divergence (F_ST_) between strains of the MAT-Prox1 and MAT-Prox2 haplotypes: D-E. Nucleotide diversity within pools of strains for each MAT-Proximal haplotype; F-G-H Tajima’s D for all strains pooled and within pools of strains for each MAT-Proximal haplotype; The MAT-proximal region defined from LD analyses and the inversion between M1400 and M6697 genomes are indicated with red and blue arrows respectively. The mating-type locus location is shown with a green triangle. The location of the putative centromere is indicated by a yellow circle. All population statistics were computed for the 1990 Swiss population along the mating-type contig (tig_0000001) of the M1400 genome per 50-kb window overlapping over 10 kbp. Windows containing fewer than 5 SNPs were removed from the analysis.

### Higher genetic differentiation and lower genetic diversity within the MAT-proximal region than in recombining regions

When analyzing polymorphism within the invasive 1990 Swiss population using the M1400 genome as reference, we found much higher genetic differentiation between strains of the two MAT-proximal haplotypes in this region than elsewhere along the genome (Fig 4C; Fig S6A; Table S4), as expected for a non-recombining region in LD with the mating-type locus. For example, the F_ST_ values between the two non-recombining haplotypes within the MAT-proximal region in the 1990 Swiss population were nearly maximal (median F_ST_=0.93 per 50 kb window), while they were near zero in the rest of the genome (median F_ST_=0.02; Wilcoxon test p-value < 2e-16 with Bonferroni correction; Fig 4C; Fig S6A). Such F_ST_ values close to 1 indicate a lack of shared polymorphism and therefore support the inference of recombination suppression in the MAT-proximal region.

Within each haplotype, the genetic diversity at the MAT-proximal region was lower than in the rest of the genome (Wilcoxon test p-value < 2e-16 for the MAT-Prox1 haplotype; p-value = 2.3e-07 for the MAT-Prox2 haplotype; Fig 4E-F; Fig S6C; Table S4), as expected for a region without recombination associated with the mating-type locus, as its effective population size is half as in recombining regions. The diversity was especially low in the pool of isolates with the MAT-Prox1 haplotype (see all MAT-Prox1 sequences clustering on a single point on the PCA in Fig 2A and in the neighbor-net network in Fig 2D, in contrast to the more scattered MAT-Prox2 sequences). Such a very low diversity may be due to a recent selective sweep or to particularly strong bottleneck in this haplotype during the invasion.

We detected signals for long-term balancing selection in the MAT-proximal haplotypes. Computation of the Tajima’s D statistics within all strains suggested signatures of balancing selection in the MAT-proximal region as expected in a region without recombination associated with the mating-type locus maintained at a frequency close to 0.5. Tajima’s D values in the MAT-proximal region pooling all sequences (median D=3.0 per 50 kb window) were higher than in the rest of the genome (median D=-0.14; Wilcoxon test p-value < 2e-16 with Bonferroni correction; Fig 4G; Fig S6D). In contrast, Tajima’s D was significantly lower than the rest of the genome within each pool of haplotype, including in the MAT-Prox2 haplotype, suggestive of positive selection (Wilcoxon test p-value = 0.0037 for the MAT-Prox1 haplotype; p-value = 0.0022 for the MAT-Prox2 haplotype with Bonferroni correction; Fig 4H-I; Fig S6D; Table S4). We found no difference in population diversity statistics between the recombining part of the mating-type contig and the other contigs (Table S4A).

The study of synonymous divergence (d_S_) between the shared orthologs in the M1400 and M6697 haplotypes suggests that their differentiation is at least 1.5 million years old and that there is no pattern of evolutionary strata within the MAT-proximal region, i.e. segments with different levels of differentiation between haplotypes that would indicate stepwise expansion of recombination suppression away from the mating-type locus. Per-gene synonymous divergence (d_S_) is typically used for detecting gradual expansion of recombination cessation and for estimating its age, as it is considered a good proxy for the time since recombination suppression. Indeed, when recombination is suppressed, mutations accumulate independently in the two non-recombining haplotypes. We plotted the per-gene synonymous divergence (d_S_) between M1400 and M6697 along the M1400 genome, as its mating-type contig is likely being assembled as a full chromosome (Fig S7A & E; Table S4). Consistent with recombination suppression, we found significantly higher d_S_ values in the MAT-proximal region (computed for 71 genes) than in the other regions of the mating-type contig (pairwise Wilcoxon test p-value <2e-16 with Bonferroni correction) and other contigs (pairwise Wilcoxon test p-value <2e-16 with Bonferroni correction). We found no significant differences between the other regions of the mating-type contig and other contigs (pairwise Wilcoxon test p-value = 0.23 with Bonferroni correction). The d_S_ pattern displayed no indication of gradual expansion of recombination cessation within the MAT-proximal region, as there was no stair-like pattern. Using synonymous substitution rate estimates across multiple ascomycetes (from 0.9 x 10^-9^ to 16.7 x 10^-9^ substitutions site per year (Kasuga et al. 2002; Taylor and Berbee 2006) and the mean d_S_ value (0.0495) across the genes shared between the MAT-proximal haplotypes, we estimated the age of their divergence to be at least 1.5 Million years (Table S4).

### Enrichment in transposable elements in the non-recombining region and inversion breakpoints

When studying the M1400 and M6697 high-quality assemblies, we detected an enrichment in transposable elements (TEs) in the MAT-proximal region in the two haplotypes compared to the rest of the genome (Fig 4B). The percentage of bp occupied by TEs (TE load) was higher than 50% in the MAT-proximal region, while it was only 9% on average in other regions. TE load in the MAT proximal region was higher in the M6697 haplotype (76%, MAT-1; MAT-Prox1) than the M1400 haplotype (68% MAT-2; MAT-Prox2). Class I retrotransposons with Gypsy (LTR-Ty3) and LARD elements were the most abundant TEs in all genomic regions (autosomes, MAT-Proximal region and the rest of the mating-type contig), representing >70% of TE annotations (Fig 5A-B), as shown previously (Demené et al. 2022). The TE load however varied significantly among TE families and genomic regions (ANOVA; Table S5). In M6697 for example, TIR elements appeared less abundant in the non-recombining than recombining regions while LTR-Ty3 elements were more frequent in the MAT-proximal region (Fig 5B). In M1400, LARD elements were more frequent in the MAT-proximal region than in recombining regions (Fig 5B).

**Figure 5:**
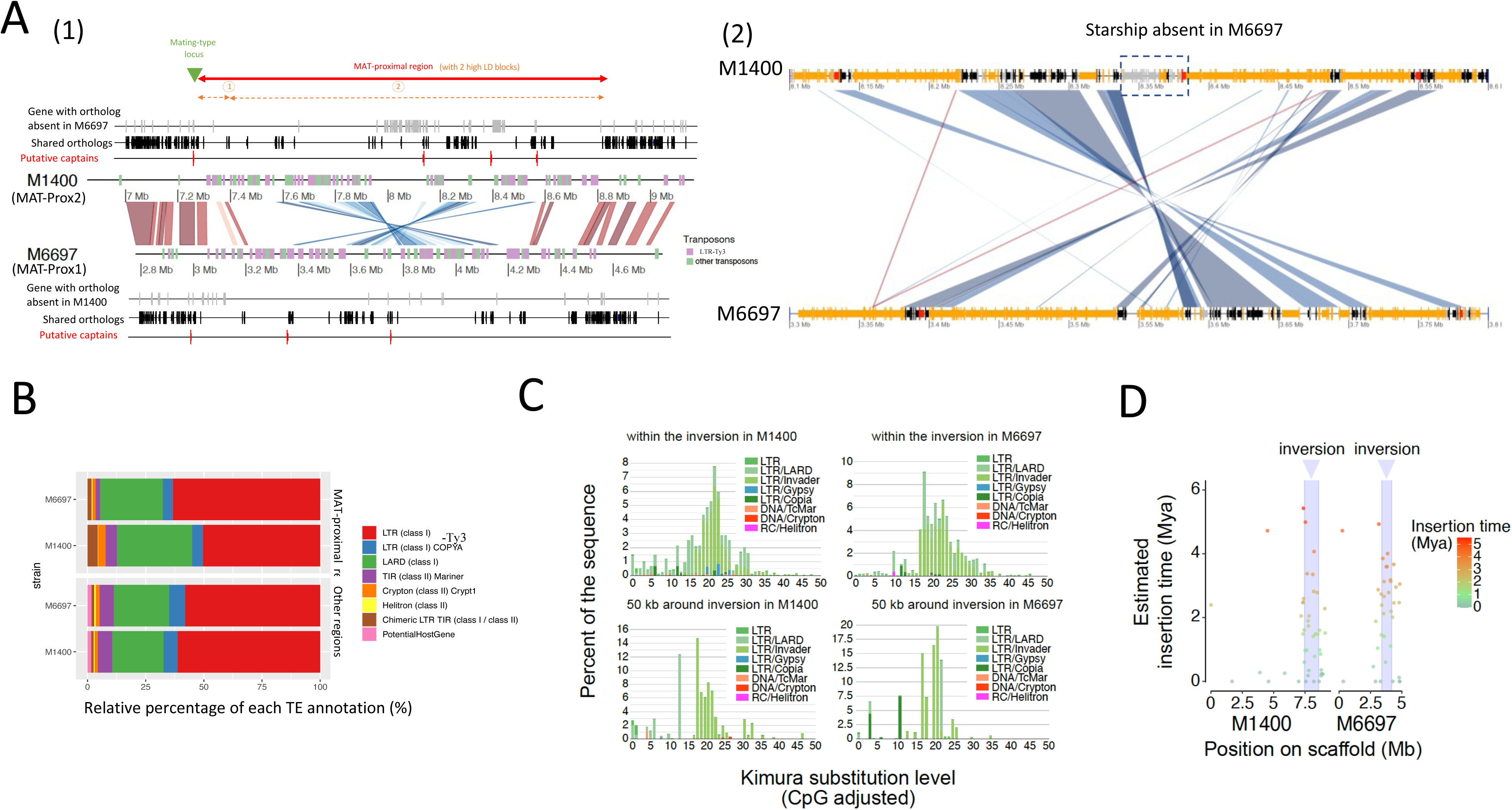
Annotation of transposable elements (TEs) and estimates of their insertion time in the MAT-proximal region lacking recombination and other genomic regions in *Cryphonectria parasitica*. A. Annotation of transposable elements (TEs) and gene density along the MAT-proximal region in M1400 and M6697 genomes. On the left panel (1) the figure shows the genomic region from 7 Mb on tig_0000001 of the M1400 genome and from 2.8 Mb on tig_0000060 of the M6697 genome. The mating-type locus is located with a green triangle. The two high-LD blocks within the MAT-proximal region are shown with orange arrows. Synteny for 10 kb segments with identity > 90 % is shown in red and inversion in blue. Transposons larger than 5 kb are shown in purple for the transposons annotated as LTR-Ty3 by and in green for other transposons. Orthologous genes shared between genomes are shown in black and unique to each each genome in grey. Genes with DU3435 domains (putative starship captains) are shown in red. On the right panel (2) the figure shows a zoom on the starship present in M1400 and absent in M6697. Transposons larger than 150 pb are shown in orange; orthologous genes shared between genomes are shown in black and unique to one genome in grey and genes with DU3435 domains are shown in red. Synteny for segments with identity > 90 % is shown in red and inversion in blue. B. Relative percentage of each TE annotation (in %) in the non-recombining MAT-proximal region and other recombining regions of M6697 and M1400 genomes. C. Pairwise genetic distance between TE copies within the inversion and around (50 kb) in both M1400 and M6697 genomes. Distribution of the Kimura substitution levels computed using the consensus sequence for the TEs. D. Estimates of the age of intact copies of LTR within and around the inversion.

The presence of transposable elements at the two inversion breakpoints in the two genomes (Fig 5A) suggests that the inversion may have occurred via non-homologous recombination mediated by these elements. By dating retrotransposon insertions using between-copy divergence and within-copy LTR divergence in both M1400 and M6697 genomes (Fig 5C and D), we found that TEs within the inversion and around the inversion breakpoints were older than TEs farther from the breakpoints. This is expected in non-recombining regions as selection is less efficient to purge TE insertions in such regions so that they remain there longer (Duhamel et al. 2023). The date estimates of LTR element insertions (Fig 5D) further indicated that the TE accumulation in the MAT-proximal region was as old as a few million years, confirming the age estimated based on sequence divergence between MAT-proximal haplotypes.

### Gene function in the MAT-proximal region and association to known phenotypes

We found no gene function known to be involved in mating or mating compatibility, nor in pathogenicity, in the MAT-proximal region (Table S7-Table S8). We nevertheless identified genes potentially coding for proteins with kinase domain or proteins with homeodomains (Table S6), that are often involved in the regulation of developmental pathways.

By re-analysing data from a previous study in the light the two haplotypes (Stauber et al. 2021; Stauber et al. 2022), we found no association of the MAT-Proximal haplotypes to previously described vegetative compatibility groups or to the sensitivity to virus infection, but more specific studies need to be performed to specifically test hypotheses.

### The MAT-proximal region harbours *Starship* elements

In addition to a high density of transposable elements, gene annotations of the MAT-proximal region in the M1400 and M6697 genome assemblies revealed multiple genes encoding tyrosine recombinases (genes with DUF3435 domains, that can be *Starship* captains) and gene content variation (Tables S6-S7), suggesting the presence of *Starships*. By running the Gene Finder Module of the starfish pipeline designed to identify *Starships* (Gluck-Thaler and Vogan 2024), we in fact identified the presence of putative *Starship* captains in the M1400 haplotype (MAT-Prox2) and three putative *Starship* captains in the M6697 haplotype (MAT-Prox1). We detected eight and ten other putative *Starship* captains elsewhere in the M1400 and M6697 genomes, respectively, but the MAT-proximal region appeared enriched in these elements (Fisher’s exact test p-value =0.0004). A phylogenetic analysis with previously described *Starship* captains (Gluck-Thaler and Vogan 2024) suggested that the captains in the MAT-Proximal region belonged to the Phoenix *Starship* family, that preferentially target AT-rich sites (Fig S7; (Gluck-Thaler and Vogan 2024)). As the Starfish pipeline performs poorly in TE-rich regions (Gluck-Thaler and Vogan 2024), we investigated manually the delimitation of putative starships. Three putative captains shared orthology relationships between the two haplotypes, one being just at the border of the MAT-proximal region near the mating-type locus (Figure 3). The high similarity between captains shared by the two haplotypes (>90% identity), their similar positions and reverse strand orientations suggested that these captains were present before the differentiation between haplotypes and before the inversion (Figure 5). The genes near these three shared captains are mostly common to the two haplotypes. The fact that these elements were present in all high-quality assemblies at the same locus prevented delimiting Starship boundaries and these genes may represent solo Starship captains or pseudogenes as found in high frequency in Pezizomycotina genomes (Gluck-Thaler and Vogan 2024). The MAT-Prox2 haplotype carried an additional putative captain, followed by 18 genes lacking in the MAT-Prox1 haplotype (Figure 5A), suggesting the presence of a *Starship* element, together with its cargo genes, only in the MAT-Prox2 haplotype. Functional annotation of the putative cargo genes included two genes with DUF3723 domain and one gene with ferredoxin reductase-type domain, that are often found in *Starships* (Table S8 (Gluck-Thaler and Vogan 2024). This putative *Starship* was about 44 kb in the M1400 strain.

In addition to the genes only present in the putative Starship specific to the MAT-Prox2 haplotype, the two haplotypes carried additional specific genes, especially the MAT-Prox2 haplotype. Based on automatic gene annotation and reconstruction of orthologous relationships, we found that, out of 175 groups of single-copy genes in the MAT-proximal region, 80 (46%) were only present in M1400 (MAT-Prox2) and 24 (14%) were only present in M6697 (MAT-Prox1). This level or presence/absence gene polymorphism was higher than in other genomic regions, in which only 3% of genes were only present in one of the two genomes genome (Fisher’s exact test p-value < 2.2e-16). However, not all genes specific to one haplotype were located close to a putative captain (Figure 3), so that part of the observed gene presence/absence polymorphism between the two MAT-Proximal haplotypes may also be due to gene losses due to degeneration or pseudogenisation after *Starship* insertions, to recombination suppression or to gene gains by other means than *Starships*.

Among the genes specific to M6697 (MAT-Prox1), we found a gene encoding a protein of the Sirtuin family (IPR003000). A Sirtuin homolog was found in a *Starship* associated with local thermal climate adaptation in the wheat fungal pathogen *Zymoseptoria tritici* (Tralamazza et al. 2024).

### Gene expression in the MAT-proximal region

We analysed the expression data available for the EP155 strain (Chun et al. 2020), carrying the MAT-Prox1 haplotype, to test whether some of the genes in the MAT-Proximal region were upregulated under certain conditions. Out of 127 genes functionally annotated in the MAT-proximal region of the EP155 genome, 50 genes were supported by RNAseq data acquired *in vitro* for this strain (Chun et al. 2020), suggesting that the region contains functionally active genes (Table S6). Among these 50 genes, nine were upregulated during barrage allorecognition of the EP155 strain with a compatible strain (Belov et al. 2021) and five were upregulated during infection of the EP155 strain with the hypovirus CHV1 (Chun et al. 2020) (Table S6), suggesting that the MAT-Proximal region may have a role in fungus-mycovirus interactions and vegetative incompatibility. Unfortunately, no expression data are available for a strain with the MAT-Prox2 haplotype, that harbours the additional *Starship* and the highest number of specific genes and no expression data is available during plant infection.

### Degeneration in the non-recombining region

The MAT-Proximal region was enriched in TEs and depleted in genes (Fig 4 and Fig 5A), which supports the inference of an old full recombination suppression. We also found overall higher non-synonymous substitution rate compared to the baseline substitution rates (d_N_/d_S_ values) between the M1400 and M6697 genomes in the MAT-proximal region (median per gene d_N_/d_S_ = 0.80) than in other contigs (d_N_/d_S_ = 0.43; pairwise Wilcoxon test p-value = 1.7e-07 with Bonferroni correction; Fig S8A; Table S4). This is consistent with relaxed purifying selection due to recombination suppression in the MAT-proximal region. In addition, 13 genes displayed d_N_/d_S_ > 1 between M1400 and M6697, suggesting that they may evolve under positive selection (Table S9).

### Recombination suppression and *Starships* in the MAT-Proximal region in other populations assessed from high-quality genome assemblies

We analyzed five additional high-quality genome assemblies for strains from the Asian native range (CL2 and CL4 clusters) and the invasion range (including two North American strains belonging to the CL1 cluster and one European strain introduced directly from Asia and belonging to the CL2 cluster), three of which were generated for the present study (Table S3). Most of the high-quality genomes displayed the same MAT-Proximal chromosomal arrangement as M1400 (MAT-Prox2, Swiss population and belonging to the CL1 cluster), suggesting that it is the ancestral state (Fig S9A-D), although the incomplete assemblies may prevent detecting other rearrangements. The inversion detected in M6697 (MAT-Prox1, Swiss population and belonging to CL1 cluster) was also present in the EP155 strain (MAT-Prox1, North America and belonging to CL1 cluster, Crouch et al, 2020; Fig S8D), indicating that this inversion was already segregating in early invasive populations in North America. In other genomic regions outside of the MAT-Proximal region, all analyzed high-quality genomes were colinear to the ESM15 reference genome, except the highly rearranged EP155 genome, as previously reported (Demené et al. 2022).

The MAT-Proximal region in all the high-quality genomes, from Asian native strains and invasive strains, displayed signs of degeneration, with a high TE load (Fig6; Fig S9), confirming the ancient age of recombination suppression and its occurrence in the native range. Study of d_S_ and d_N_/d_S_ between the MRC10 strain, belonging to the European clonal lineages of direct Asian origin (CL2 cluster), the M6697, M1400 and the two Asian native strains further confirmed recombination suppression in the native range and in invasive strains from different origins (North America or directly from Asia). The d_S_ and d_N_/d_S_ values were indeed significantly higher in the MAT-proximal region than other genomic regions (Fig S7 B-E). Mean d_S_ values in the MAT-proximal region were in the same range as d_S_ values between the M6697 and M1400 genomes, suggesting a similar date of recombination suppression and, therefore, that recombination suppression was ancestral. The MAT-proximal region appeared smaller in the two Asian strains XIM9508 and ESM15 (<1 Mb) than in invasive strains (Table S3), but this may be due to the incomplete genome assemblies, and/or to additional *Starship* presence/absence polymorphism.

We detected the three captains shared between the M6697 and M1400 haplotypes in the MAT-proximal region of all high-quality genome assemblies, indicating that their insertions in the MAT-proximal region were old, being already present in the native range. We found the putative *Starship* specific to M1400 (MAT-Prox2), i.e., the captain and the 18 cargo genes, in the MAT-proximal region of DUM-005 (MAT-Prox2) and in the haplotypes of MRC10 and the Japanese strain ESM15 (both genetically close to MAT-Prox2; Figure S10D1), while it was absent in the MAT-Proximal region of the EP155 strain (MAT-Prox1). In the XIM9508 Chinese strain (genetically close to MAT-Prox1; Figure S10D1), the captain and four genes of the *Starship* specific to M1400 were absent in the MAT-proximal region. However, 14 of its putative cargo genes were present and a very similar captain was found elsewhere in the genome, suggesting that this *Starship* may have moved away from the MAT-Proximal region after its insertion there. In the seven high-quality assemblies, the putative *Starship* presence/absence polymorphism detected between the 1400 and M6697 genomes thus seem associated to the MAT-Proximal haplotypes in the invaded range and also in the native Asian range (Figure S10D), although this would need to be confirmed with additional high-quality assemblies.

### Presence of the two MAT-Proximal haplotypes in other populations, but less differentiated in the native range

By analyzing available additional Illumina genomes of invasive strains from the first invasion wave via North America (strains from the US and European clonal lineages), we recovered the two MAT-proximal haplotypes in balanced frequencies. We indeed observed high differentiation between two clusters in the MAT-Proximal region, contrasting with a lack of genetic subdivision in other genomic regions or a completely different genetic subdivision, corresponding to previously described population genetic structures (Figs S10A and S10C). This reinforces the view of balancing selection on the MAT-Proximal haplotypes.

In the native range, the MAT-Proximal region also seemed to have undergone a very different evolutionary history than the rest of the genome (Figure 7). In particular, the neighbor-net network with multiple strains sequenced previously from the CL2 and CL3 clusters (from the invaded range and the native range, in South Korean and Japan) and from the CL4 cluster (from the native range, in China) indicated that sequences from these three clusters were intermingled in the MAT-proximal region (Fig 7A), while the structure in the rest of the genome instead corresponded to the different population clusters, i.e., CL1, CL2, CL3 and CL4 (Fig 7B). However, the two haplotypes were much less differentiated in the native range and in invasive populations originating directly from Asia than in invasive populations from the first invasion wave via North America. For example, branch lengths on the neighbor-net indicate a lower differentiation between haplotypes within CL2 (Fig 7A) than within the European CL1 or Swiss invasive populations (Fig 2D; Fig S3) and some reticulations were present between MAT-proximal haplotypes.

**Figure 6:**
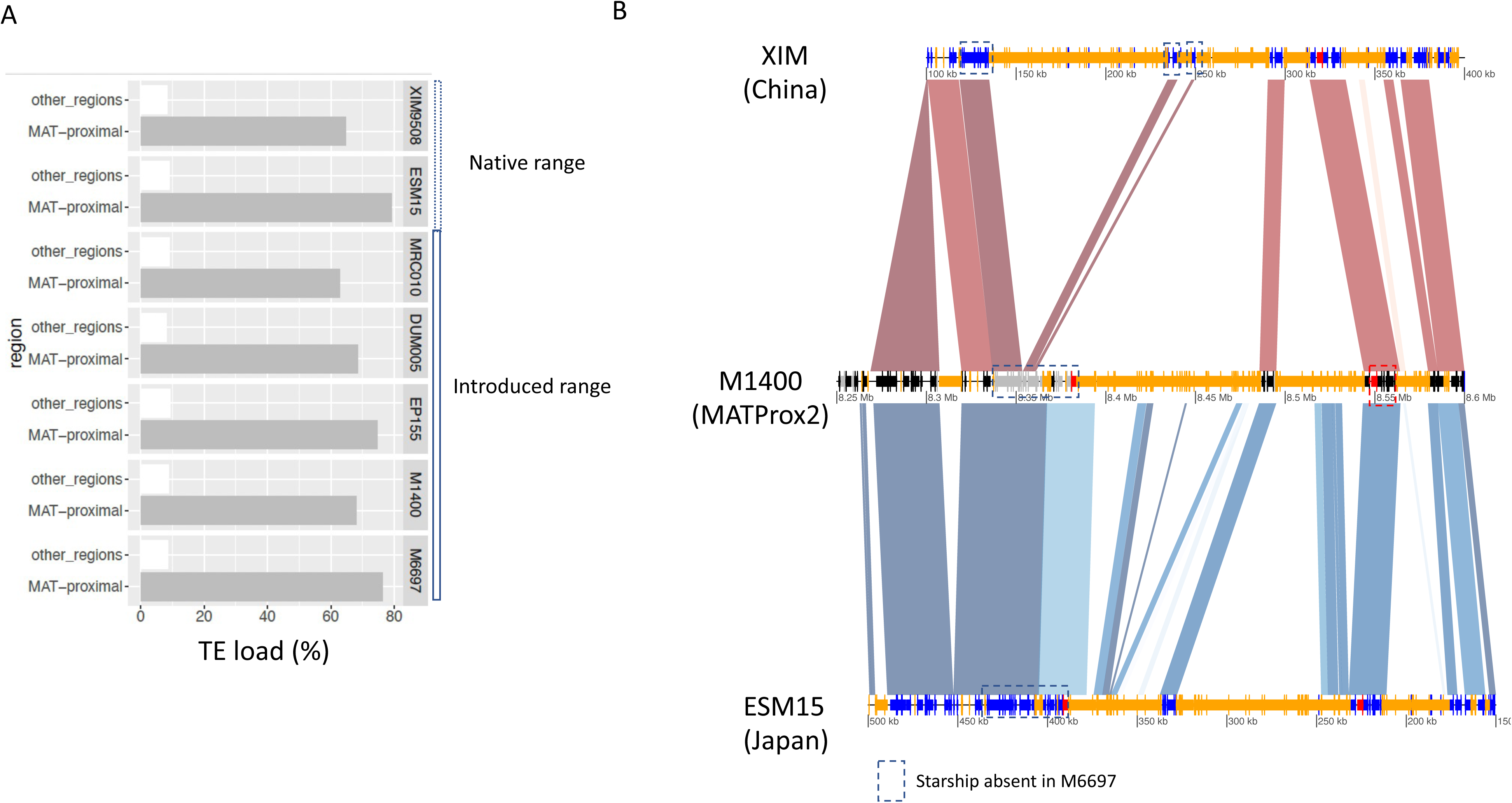
Transposable elements content and Starship content in high quality genome assemblies from other *Cryphonectria parasitica* populations from the native and the introduced range. A. TE load (percentage of base pairs occupied by transposable elements) in the seven high quality genome assemblies in the MAT proximal region and other recombining regions. **B.** Annotation of transposable elements (TEs) and gene density along the MAT-proximal region in M1400 and the two Asian genomes ESM15 and XIM9508. Orthologous genes shared between M1400 and M6697 genomes are shown in black and unique to M1400 in grey as in Figure 5. Genes of ESM15 and XIM9508 are shown in darkblue? Genes with DU3435 domains (putative starship captains) are shown in red. Transposons larger than 150 pb are shown in orange. Synteny for segments with identity > 90 % is shown in red and inversion in blue.

**Figure 7:**
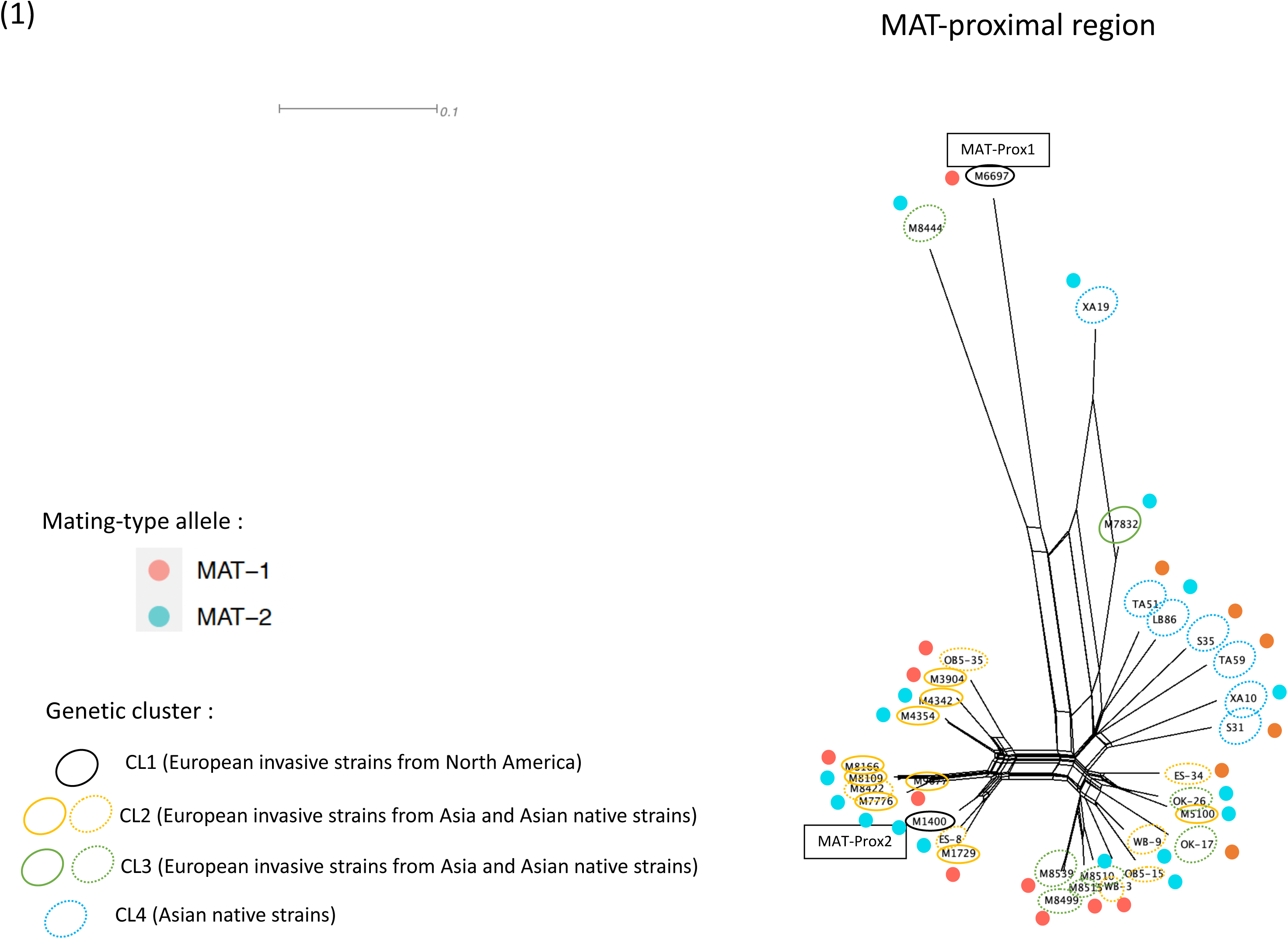
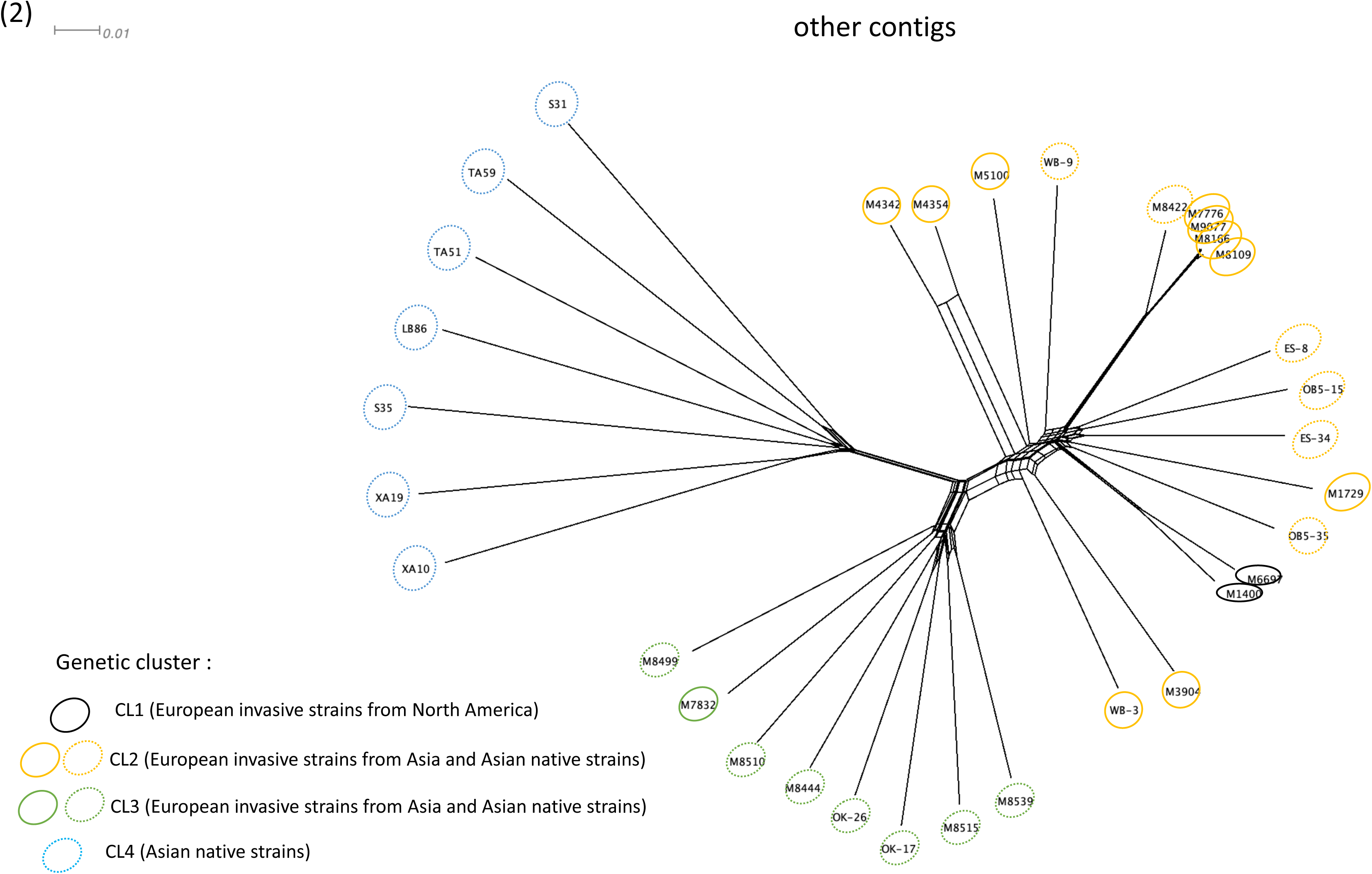
Genetic structure using single nucleotide polymorphisms (SNPs) in the MAT-proximal region lacking recombination (A) and other genomic regions (B) in other resequenced *Cryphonectria parasitica* populations from the native and the introduced range. Neighbor-net network from a SplitsTree analysis based on: **A** SNPs (n=4,120) located within the MAT-proximal region along the contig carrying the mating-type locus (scaffold_2 of the EP155 genome); **B** SNPs (n= 103,058) located on other contigs of the EP155 genome. Color of empty circles around strain ID indicate the genetic clusters the strains belong to (CL1, CL2, CL3, CL4; n= 33 strains). Dotted circles indicate strains of the native range and plain circles indicate strains of the introduced range. Red and blue dots near strain ID indicate the mating type (*MAT-1* or *MAT-2*, respectively).

We found no association of the MAT-Proximal haplotypes with mating types at the world scale (Figs 7 and S10). In the S12 European invasive clonal lineage (only MAT-1), we found a single MAT-Proximal haplotype (MAT-Prox1; Fig S10B), but we detected the two MAT-Proximal haplotypes in all other populations. Even in two invasive European lineages of American origin with a predominantly clonal structure (re092 and re103), the strains with a mating type different from their clone-mates, due to localized introgression from other clonal lineages (Demené et al. 2019), also had their MAT-Proximal region introgressed (Fig S10C); this finding reinforces the view of the strong linkage between the mating-type locus and the MAT-Proximal region, of an advantage of maintaining the two MAT-Proximal haplotypes in populations, and perhaps of maintaining an association between MAT-proximal haplotypes and mating types within populations. Three 2019 Swiss strains exhibited a similar intermediate sequence in the MAT-Proximal region as the other strains from Europe with a direct Asian origin described above (Fig S10C). This intermediate haplotype thus seemed associated with the second invasion in Europe, directly from Asia, and carried the same captains as the MAT-Prox2 haplotype from the M1400 strain. The location as an intermediate in the neighbor-net network may be due to recombination between the haplotypes, or may just reflect the fact that the two haplotypes are less differentiated in the native Asian area.

### No evidence of introgression in the MAT-proximal region from *closely related* Cryphonectria species

The stronger differentiation in the MAT-Proximal region in the Swiss and CL1 invasive populations than in other populations could also be due to an introgression event from a closely related species; however, the small genetic distance and the reticulations observed in the neighbor-net network between the M6697 Swiss reference genome and the Asian genomes M8444 and XA19 rather indicate that the MAT-Prox1 haplotype in Switzerland originated from Asian *C. parasitica* populations, likely a population different from those at the origin of the CL2 invasive cluster (Fig 7).

Gene genealogies also provided no evidence of introgression in the MAT-proximal region from closely related *Cryphonectria* species. We performed a gene orthology analysis including three genomes of the closely related species *C. japonica*, C. *carpinicola*, and *C. naterciae* that were previously sequenced with short read technologies (Stauber et al. 2020). We retrieved only nine genes from the MAT proximal region that were shared between the M6697 and M1400 genomes that had orthologs in one of the related species. This may be due to low quality assemblies in the outgroups (Stauber et al. 2020) and/or to many gene gains in this region in *C. parasitica* because of *Starships*. We found no gene with a topology consistent with introgression from another species into the MAT-Proximal region, i.e. with a MAT-Prox1 or MAT-Prox2 haplotype that would branch with an outgroup allele rather than with the alternative haplotype of *C. parasitica*. The high load of TEs and gene disruptions in the two MAT-Proximal haplotypes does not fit with the introgression hypothesis either, unless the introgression occurred from a species with recombination suppression in the MAT-Proximal region. The Starships may have moved from a distant species by horizontal gene transfer together with their cargo genes, but we found no evidence for this by blasting the captains and cargo genes in public databases.

## Discussion

### A large non-recombining region near the mating-type locus, with two highly differentiated haplotypes maintained polymorphic in the native and introduced ranges

We found strong evidence for the complete cessation of recombination in a region of more than 1 Mb (1.2 Mb to 1.5 Mb depending on the haplotype), proximal to the mating-type locus in *C. parasitica*. Indeed, we detected maximal levels of linkage disequilibrium in otherwise recombining populations and the existence of two differentiated haplotypes, without shared polymorphism, while even low rates of recombination can homogenize alleles and prevent LD build-up (Dufresnes et al. 2015). The full cessation of recombination was further supported by the existence of a previously unknown, large inversion in the center of the non-recombining region in invasive strains. This is in agreement with previous genetic maps reporting lack of recombination near the mating-type locus in crosses involving Japanese, North American and Italian isolates (Kubisiak and Milgroom 2006a). The higher TE load in this region constitutes additional evidence for full recombination cessation, as the lack of recombination renders selection against TE accumulation less efficient, in particular due to Muller’s ratchet: two chromosomes with different TE insertions cannot recombine to produce a chromosome free of TE insertions. TE takes hundreds of thousands of years to accumulate in non-recombining regions (Duhamel et al. 2023), so that the high TE load in the MAT-Proximal region in *C. parasitica* indicates old recombination cessation. The particularly high frequency of non-synonymous substitutions in the MAT-Proximal region worldwide further indicates degeneration and ancient recombination suppression. The existence of two differentiated haplotypes appeared less clear in the native range in Asia and in the second-wave invasive populations originating directly from Asia than in invasive populations from the first introduction via North America, but the MAT-Proximal region nevertheless appeared to display a different evolutionary history than the rest of the genome and a high TE load.

The estimates for the differentiation between the two MAT-Proximal haplotypes based on synonymous divergence between shared single-copy orthologs and based on TE insertion dates were at least 1.5 million years. The suppression of recombination in *C. parasitica* may be younger than this estimate if one of the two haplotypes was introgressed from a distant species. The evolution of non-recombining regions by introgression has been reported for example in ants and butterflies (Jay et al. 2018; Helleu et al. 2022; Stolle et al. 2022). However, we found little evidence for an introgression: i) the two haplotypes were detected in native populations and in the CL2 cluster of invasive European populations directly originating from Asia, even if less differentiated than in the invasive populations from the first introduction, ii) the MAT-Prox1 haplotype in the CL1 cluster that is the most differentiated from the MAT-Prox2 haplotype is genetically close to an Asian haplotype, iii) footprints of recombination suppression are present in the native range in terms of degeneration and particular genomic structure in the MAT-Proximal region, and iv) we found no signatures of introgression from closely related species in gene genealogies or by blast in public databases. In any case, the findings of recombination suppression footprints and of the presence of two differentiated haplotypes in the two invaded continents, Europe and North America, as well as in the native range of *C. parasitica*, with intermingled sequences from different Asian populations in neighbor-net networks, indicate ancient recombination suppression and long-term polymorphism maintenance in the MAT-Proximal region. These findings point to a strong balancing selection maintaining two differentiated haplotypes in *C. parasitica*.

In contrast to other known non-recombining regions on sex or mating-type chromosomes (Bergero and Charlesworth 2009; Hartmann et al. 2021), the region without recombination was not completely linked to the mating-type locus. Indeed, the occurrence of rare recombination events between the non-recombining MAT-Proximal region and the mating-type locus are supported by previous segregation analyses (Kubisiak and Milgroom 2006b) and the lack of association found here between the MAT-Proximal haplotypes and the mating types at the worldwide scale, as well as their incomplete association within populations.

### Proximal cause of recombination suppression

The large inversion in the MAT-Proximal region may contribute to recombination suppression. There may also be additional nested inversions and/or translocations in the MAT-Proximal region as there are breaks of synteny between the inverted fragments with many transposable element insertions. The chromosomal arrangement of the M1400 MAT-Prox2 haplotype is likely the ancestral state as it is the most frequent worldwide, although a more complete sampling in the native range is required to obtain definitive conclusion. The non-recombining region was defined based on LD pattern and was larger than the inversion. This may indicate that additional proximal mechanisms suppress recombination. The inversion may even be a consequence rather than a cause of recombination cessation, as previously found in other fungal mating-type chromosomes (Grognet et al. 2014; Sun et al. 2017; Branco et al. 2018; Vittorelli et al. 2022), especially as the inversion was not detected so far in the native range despite the presence of footprints of recombination suppression. Other proximal causes of recombination cessation may for example be genetic recombination modifiers, epigenetic marks, histone modifications or lack of recruitment of proteins responsible for double-strand breaks (Maloisel and Rossignol 1998; Boideau et al. 2022; Legrand et al. 2024). The lack of rearrangements at the edge of the MAT-proximal region may allow rare recombination or gene conversion events, which may explain the presence of two distinct LD blocks with lower LD between them and the lower differentiation at the edges of the MAT-proximal region. Alternatively, the LD around the inversion may be formed by less frequent recombination as recombination is often modified around inversion breakpoints (Pegueroles et al. 2010; Stevison et al. 2011) and the actual size of the non-recombining region may be smaller than suggested from the LD pattern.

### The MAT-proximal haplotypes carry putative *Starship* elements

We detected signatures of *Starship* elements in the non-recombining MAT-proximal region, with likely three “solo” captains shared between the two haplotypes, but also an additional captain associated to 18 specific genes, strongly suggesting the presence of a cargo-mobilizing *Starship*, only present in the MAT-Prox2 haplotype. The difference in gene content between the two MAT-proximal haplotypes thus likely at least partly result from specific cargo genes inserted with *Starships,* although there could also be gene losses because of the less efficient selection due to recombination suppression, and/or additional gene gains. One of the captains shared between haplotypes was located just nearby the mating-type locus and at the edge of the MAT-proximal region. This altogether strongly suggest a role of *Starship* elements in the formation of the MAT-proximal haplotypes and in their long-term maintenance, although experiments are required to confirm this hypothesis and elucidate their role. High-quality assemblies suggested that the presence/absence of the polymorphic *Starship* insertion was associated to the MAT-Proximal haplotypes, although this also needs to be checked in a larger set of strains. The presence of the inversion in the native range also remains to be investigated. Additional sampling in the native range and high-quality genome assemblies are required to obtain a more comprehensive view of the history of the MAT-Proximal region and of the putative *Starship* insertions.

*Starships* may have been introgressed from another species, as frequent with these elements (Ropars et al. 2015; Peck et al. 2023), but searches in databases returned no significant blast results. Such horizontal gene transfers should not have, however, inflated the date estimates of divergence of recombination suppression in the MAT-Proximal region, as these were computed based on shared single-copy orthologs between haplotypes and the shared putative *Starship* elements seem to correspond to homologous insertions.

### The selective forces potentially maintaining the two haplotypes in the MAT-proximal region

Our findings indicate that selection maintains the two haplotypes polymorphic in the MAT-Proximal region. There can be several hypotheses regarding the nature of this balancing selection. The degeneration of the non-recombining region may help maintaining the two haplotypes by selection due to the sheltering of combinations of deleterious mutations in repulsion, i.e. thanks to a heterozygote advantage called pseudo-overdominance (Abu-Awad and Waller 2023). Such pseudo-overdominance advantage may only be a consequence of recombination suppression, and contribute to its maintenance, or may even be its initial cause (Branco et al. 2017; Jay et al. 2021; Charlesworth 2023; Jay, D.L. Jeffries, Hartmann, et al. 2024). These hypotheses require the existence of a substantial diploid or dikaryotic phase for sheltering recessive deleterious alleles in a heterozygous state. In fact, although *C. parasitica* has a predominant haploid phase, strains can be found as dikaryons heterozygous for the mating-type locus in some natural populations, and not only as monokaryons (McGuire et al. 2004; McGuire et al. 2005; Stauber et al. 2021; Stauber et al. 2022). The importance and frequency of heterokaryons in *C. parasitica* in nature would deserve further investigations. It may be sufficient if recessive deleterious alleles are sheltered in a substantial percentage of individuals for selecting recombination suppression, and/or if the genes involved are preferentially expressed during the diploid or dikaryotic phases. This hypothesis of balancing selection, and in particular the hypothesis of overdominance (i.e. heterozygote advantage), does not require the same association between the MAT-Proximal haplotypes and mating types across all populations worldwide, it is sufficient that the MAT-Proximal haplotypes are strongly associated to mating types within populations. Further sampling will be required to test whether the MAT-Proximal haplotypes are associated to mating types within local populations in the CL3 and CL4 clusters and in the native Asian range. It will also be interesting in future studies to compare the fitness between strains carrying the two MAT-Proximal haplotypes and dikaryotic strains being homozygous for the MAT-Proximal haplotype. Such experiments in Sordariales fungi reported a heterozygote advantage (Guyot et al. 2024). Results from previous studies suggest that strains carrying either the same or different MAT-Proximal haplotypes can be crossed in *C. parasitica* (Stauber et al. 2021).

It could indeed also be that genes evolve under overdominance in this region for other reasons than deleterious mutations, so that a partial linkage to a permanently heterozygous locus is beneficial, perhaps in relationship with the pathogenic lifestyle of the fungus. One could imagine, for example, that heterozygosity could be advantageous at genes involved in virulence against the tree host, and especially the novel hosts colonized in invasive ranges, or in resistance against a parasitic virus known to negatively affect fitness in *C. parasitica* (Choi and Nuss 1992; Brusini et al. 2017). As a matter of fact, multiple genes present in the MAT-Proximal region are up-regulated under infection by the hypovirus CHV1 or during the vegetative incompatibility reaction considered to play a role in the prevention of virus transmission (Choi et al. 2012; Rigling and Prospero 2018). The specific genes unique to one or the other MAT-proximal haplotypes, and possibly the cargo genes inserted with the *Starships*, may contribute to such a heterozygous advantage.

As an alternative to a heterozygote advantage, the selection maintaining the recombination suppression and the two haplotypes in the MAT-proximal region could be some kind of negative-frequency dependent selection of beneficial allelic combinations, possibly linked to a trench-warfare-like arms race with the host tree, the virus or the microbial community, or to self-incompatibility, where partial linkage to the mating-type locus would help maintenance in balanced proportions and therefore would prevent allele loss (Tellier et al. 2014; Jay, Aubier, et al. 2024). The MAT-proximal region did not include any of the genes previously identified as controlling vegetative incompatibility in *C. parasitica* but not all self-incompatibility genes have been identified yet. The MAT-proximal region actually carried genes upregulated under virus infection or vegetative incompatibility reaction. Such a role in host-pathogen interactions or vegetative incompatibility would also be consistent with the selective sweep footprints detected in the MAT-proximal region in a previous study (Stauber et al. 2021), if there is recurrent positive selection for improving the efficiency of the pathogen weapons within each of one of the two MAT-proximal haplotypes or new, rare self-recognition alleles. Such negative-frequency dependent selection, could also explain the particular population structure in the MAT-proximal region in the native region, with a mix of sequences from the different genetic clusters, if alleles introgressed between populations are favored by a positive selection of rare alleles, or with the long-term maintenance of self-incompatibility alleles, and therefore of ancestral polymorphism. Much higher differentiation between MAT-Proximal haplotypes in the invasive than the native populations may be due to a selective sweep of a rare and differentiated MAT-Prox1 allele present in the native range or having evolved rapidly, by particular demographic effect during the invasion (Moinet et al. 2022), both of which are consistent with the very low genetic diversity in the MAT-Prox1 haplotype in the introduce range. Balancing selection and location adjacent to a non-recombining region has been reported for loci involved in host resistance in the *Daphnia– Pasteuria* system (Fredericksen et al. 2023). The MAT-Proximal region may in this case even include genes involved different traits under balancing selection, and would then constitute a supergene (Schwander et al. 2014). The MAT-Proximal region also contained, only in the MAT-Prox1 haplotype, a gene of the Sirtuin family, a homolog of which had previously been found in a *Starship* associated with local thermal climate adaptation in the wheat fungal pathogen *Zymoseptoria tritici* (Tralamazza et al. 2024). Another hypothesis to explain the balancing selection in the MAT-Proximal region may thus be a heterogeneous selection, with different genes in the two haplotypes providing contrasted advantages in different conditions or different phases of the life cycle. The proximity of the mating-type locus, permanently heterozygous in dikaryotic and diploid stages, may help maintaining balanced frequencies of the two MAT-proximal haplotypes. Here too, the genes inserted by the *Starships* may play a role in such balancing selection, especially the genes that are present in a single of the two MAT-Proximal haplotypes. The other transposable elements insertion polymorphism observed from the high quality assemblies may also be adaptive (Casacuberta and González 2013; Orteu et al. 2024).

Another hypothesis for explaining such extension of recombination suppression beyond the mating-type locus is antagonistic selection, i.e. linkage of alleles that would improve fitness of a MAT-1 gamete while being deleterious in a MAT-2 gamete, or vice-versa. However, full recombination suppression with the mating-type locus would be expected under such antagonistic selection, as well as the same association between the MAT-Proximal alleles and the mating-type in all populations, in contrast to our findings. In addition, we found no particular predicted function in the MAT-proximal region that could be related to mating compatibility and there are very little functions, if any, with possible antagonistic functions between mating types in fungi (Bazzicalupo et al. 2019; Hartmann et al. 2021).

### Conclusion

In conclusion, we provide strong evidence for the existence of a non-recombining region partially linked to the mating-type locus in the chestnut blight fungus *C. parasitica*, with two highly differentiated haplotypes, each carrying specific genes, maintained polymorphic by selection. We found footprints of balancing selection in the MAT-proximal region in the first introduction of the pathogen in Europe from North America and a chromosomal inversion. The non-recombining region also displayed footprints of particular evolution in Asia and in the second invasion wave directly from Asia, although the levels of differentiation between haplotypes was lower than in the populations from the first introduction wave. This non-recombining region may underlie important adaptive traits and thereby provide important applications for the control of a devastating tree pathogen. This is supported by the finding of putative *Starships* elements in the MAT-proximal region, i.e., giant mobile elements recently discovered in ascomycete fungi, containing multiple cargo genes (Gluck-Thaler et al. 2022; Urquhart et al. 2024), that can be involved in adaptation. In addition, the high-quality genome assemblies provided here, from the native and invaded ranges, will more generally be useful for studies aiming at understanding the evolution of this invasive and damaging pathogen.

## Material and Methods

### Strains and genomic data

For population genomic analyses, we studied a collection of 386 monokaryotic *C. parasitica* strains sampled worldwide, from the native and invaded ranges of the pathogen, and sequenced previously with the short-read Illumina technology (strain information are presented in Table S1; (Demené et al. 2019; Stauber et al. 2021; Stauber et al. 2022)). Data were downloaded from NCBI Bioproject numbers PRJNA604575, PRJNA644891 and PRJNA706885. We focused our analyses first on European invasive strains originating from North America. We studied 88 strains belonging to the CL1 genetic cluster, in central and southeastern Europe (Stauber et al. 2021). We excluded the putative clonal genotypes previously identified (Stauber et al. 2021). We also studied 71 strains sampled in the 1990 and 62 strains sampled in 2019 in southern Switzerland (Stauber et al. 2022). Mating-type ratios close to ½ and population structure analyses of the CL1 genetic cluster and the Swiss 1990 population suggest frequent recombination in these populations (Stauber et al. 2021; Stauber et al. 2022). The mating-type ratio was 33% *MAT-1* in the 2019 Swiss population and population structure analyses suggested regular sexual reproduction and recent population bottleneck. To study the presence of the two haplotypes in the MAT-proximal region more broadly, we analyzed monokaryotic strains belonging to the CL2, CL3, CL4 genetic clusters, as well as the S12 European invasive lineage (Stauber et al. 2021) and additional monokaryotic strains from the US and Europe (with an Asian or North American origin (Demené et al. 2019)). We excluded heterokaryotic strains, i.e. strains having both MAT-1 and MAT-2 alleles, as the phase was challenging to infer.

For mapping and SNP calling, we first used as reference the 43.9 Mb genome sequence EP155 v2.0 of the strain EP155 (*MAT-2,* North America, CL1 cluster) available at the Joint Genome Institute (http://jgi.doe.gov/)(Crouch et al. 2020). For comparative genomics, we used the published genome of the strain ESM15 (MAT-2, Japan, CL2 cluster; (Demené et al. 2022)) available at DDBJ/ENA/GenBank on the bioproject PRJNA700518 under the accession JAGDFO000000000. We additionally sequenced *de novo,* with long-read technologies, five strains from the native and invaded ranges of the pathogen. We sequenced with PacBio Hifi the genomes of the strains M1400 (MAT-2) and M6697 (MAT-1) sampled in Gnosca, southern Switzerland (Stauber et al. 2021). Mycelia were stored as glycerol stocks at -80 C after strain isolation. To produce mycelium for DNA extraction, isolates were inoculated onto cellophane-covered potato dextrose agar plates (PDA, 39 g/L; BD Becton, Dickinson and Company, Franklin Lakes, USA) (Hoegger et al. 2000) and incubated for a minimum of 1 week at 24°C, at a 14 hr light and 10 hr darkness cycle. Mycelium and spores were harvested by scratching the mycelial mass off the cellophane, transferring it into 2 mL tubes and freeze-drying it for 24 hr (Stauber et al. 2021). DNA extraction was performed with the NucleoBond High Molecular Weight DNA kit from Macherey-Nagel, with the mechanical disruption of about 30 mg of lyophilized mycelium with two beads of 3 mm for 5 min at 30 Hz. Sequencing was outsourced to Novogene, the Netherlands. We additionally sequenced with Oxford Nanopore MinION technology the strains XIM9508 (China, CL4 cluster, MAT-1), MRC10 (South Western France introduced directly from Asia, CL2 cluster, MAT-2) and DUM-005 (USA, CL1 cluster, MAT-2). These isolates had been collected for previous studies (Milgroom et al. 1996; Liu et al. 2003; Dutech et al. 2012). The protocols for mycelium production, DNA extraction and sequencing for these three strains were the same as in (Demené et al. 2022).

### Short-read data processing and SNP calling

We used SNP calling datasets of the genomes from monokaryotic strains of the CL1, CL2, CL3, CL4 genetic clusters, the S12 invasive lineage and the Swiss populations against the EP155 reference v2.0 genome obtained in (Stauber et al. 2021; Stauber et al. 2022). We performed mapping and raw SNP calling of short-read data using the M1400 new genome assembly as a reference as described in (Stauber et al. 2021; Stauber et al. 2022). Briefly, we trimmed reads with Trimmomatic v0.36 (Bolger et al. 2014) and aligned them with Bowtie 2 v2.3.5.1 (Langmead et al. 2009) and SAMtools v1.9 (Li et al. 2009) to the EP155 v2.0 genome. Raw SNP calling and filtration for quality were conducted with the genome analysis toolkit GATK v3.8 and v4.0.2.0 (McKenna et al. 2010). We used the filtration parameters described in (Stauber et al. 2021): QUAL>=100, MQRankSum (lower) >= -2.0, QD:_20.0, MQRankSum (upper) <=2.0, MQ:_20.0, BaseQRankSum (lower) >=-2.0, ReadPosRankSum (lower) >=-2.0, ReadPosRankSum (upper) <=2.0, BaseQRankSum (upper) <=2.0. We further removed SNPs overlapping with transposable elements predicted *de novo* in the EP155 v2.0 genome by (Stauber et al. 2021).

### Population genomics analyses

For all population genomics analyses, we excluded SNPs with missing data in more than 10% of the strains and kept only polymorphic strains with vcftools v0.1.16 (Danecek et al. 2011). To study linkage disequilibrium, we further excluded rare variants (minor allele frequency <0.1) with the vcftools (Danecek et al. 2011) option --maf 0.1. We computed LD with the -- hap-r2 option of vcftools (Danecek et al. 2011) for each scaffold and each population separately. We used the --thin 50000 option of vcftools (Danecek et al. 2011) to sample SNPs distant of at least 50 kb. We used the R package LDheatmap v1.0-6 (Shin et al. 2006) to plot LD r^2^ values among SNP pairs. To perform principal component analyses (PCAs), we first used vcftools (Danecek et al. 2011) to convert VCF format files in Plink format. We then used the Plink v1.90b5.3 (Purcell et al. 2007) –pca command to run PCA analysis. We used the R package PopGenome v2.7.5 (Pfeifer et al. 2014) to compute nucleotide diversity, Tajima’s D values and the F_ST_ index in 50 kb window overlapping over 10 kbp. Windows containing fewer than 5 SNPs were removed from the analysis. We used the R package ggplot2 (2_3.5.0) to plot results. Pairwise Wilcoxon tests were performed in R with Bonferroni or false discovery rate correction.

We built neighbornet networks using SplitsTree4 v 4.19.2 (Huson 1998). VCF file were converted to nexus format using PGDSpider v1.0 tool (Lischer and Excoffier 2012). For the study of other populations, we included only one strain of each haplotype of the invasive European population but not all CL1 and Swiss strains for network readability; we nevertheless checked that the results were the same with all CL1 and Swiss strains.

### Long-read based assemblies

PacBio Hifi reads of strains M1400 and M6697 were both assembled using canu v1.8 (Koren et al. 2017) program with a set genome size of 44 Mb. Multiple assembly pipelines were used for the other strains. Oxford Nanopore MinION reads and Illumina reads of the genomes of both XIM9508 and MRC10 strains were assembled using HybridSPAdes (Antipov et al. 2016). Assemblies were manually curated and scaffolds were cut when an evidence of a chimeric connection was detected (i.e. mis-paired short reads) as previously described (Demené et al. 2022). For DUM005, the assembly was generated by Ra with basic parameters that uses Oxford Nanopore MinION reads and corrects the assembly with Illumina reads (https://github.com/lbcb-sci/ra) as it outperformed the HybridSPAdes assembly. As the HybridSpades assembly of the MRC10 strain suggested absence of collinearity with the MAT-proximal region M1400 and M6697, we further checked the assembly of the MAT-proximal region of MRC010 by generating a meta-assembly of this strain. We used the assembler tool canu v2.2 with an estimated genome size of 42 Mb. We also used Flye v2.9.3-b1797 (Kolmogorov et al. 2019) with an estimated genome size of 42 Mb, --nano-raw for reads with an error rate below 20 and a coverage for initial disjointig assembly of 50. Then we used ragtag (Alonge et al. 2019) to patch the canu assembly with the Flye assembly as query in a first loop. In a second loop, the canu assembly was patched with the first loop ragtag assembly. Finally, we polished this second output assembly of ragtag with short reads and the consensus part of medaka v1.11.3 with no change in the parameters (https://github.com/nanoporetech/medaka). Assemblies statistics were obtained with quast v5.1 (Gurevich et al. 2013). We assessed the completeness of each assembly using the Benchmarking of Universal Single-Copy Orthologue (BUSCO) tools with the Sordariomyceta ortholog set (sordariomycetes_odb10, 2020-08-05, n = 3817 orthologous groups searched) (Manni et al. 2021). Gene models were predicted with the Helixer v0.3.1 pipeline (Holst et al. 2023). We also run Helixer pipeline for ESM15 and EP155 strains for the gene orthology analysis. Statistics of the obtained gene annotation was obtained with the Agat v1.0.0 tool (Dainat et al. 2020). Transposable elements of all genomes were annotated using Repeatmasker v4-0-7 (Smit et al. 2013) and the customized library built for *C. parasitica* in (Demené et al. 2022) contained in the “Curated_TEs_database_ESM015_EP155.fa” file (available on the “Portail Data INRAE: Chromosomal rearrangements but no change of genes and transposable elements repertoires in an invasive forest-pathogenic fungus" at https://doi.org/10.15454/UTIB8U). The class Gipsy invader was renamed LTR-Ty3. We filtered out TE copies shorter than 100bp. We used the HybridSPAdes preliminary assembly of MRC10 for gene model annotations. To study gene functions of the EP155 genome, we used the gene annotation available at http://jgi.doe.gov/ (Crouch et al. 2020). To study support from RNAseq data, we used RNAseq data from the EP155 strain cultivated *in vitro* available at Genebank under Project ID number PRJNA588887 and accessions numbers SRR10428542, SRR10428543, SRR10428544 (Chun et al. 2020). Raw reads were mapped using STAR v2.7.10a (Dobin et al. 2013) with the settings. --alignIntronMax 1000 --limitBAMsortRAM 1629219965--quantMode GeneCounts. We used the program featureCounts (Liao et al. 2013) v2.0.6 with the options -p --countReadPairs -M - -B -O --largestOverlap. We considered a gene to be supported for read count >10. To predict gene functions of the protein predicted by Helixer, we used the funannotate pipeline with default options. To look for *Starships*, we ran the Gene Finder Module of Starfish pipeline with default options (Gluck-Thaler and Vogan 2024). To predict gene functions of the protein predicted by Helixer, we used the funannotate pipeline with default options. To identify putative *Starships*, we ran the Starfish pipeline with default options (Gluck-Thaler and Vogan 2024). We studied the phylogenetic relationships of the putative Starships captains by aligning them to the YRsuperfamRefs.faa from the starfish database (Gluck-Thaler and Vogan 2024). Protein sequences were aligned using Clustal Omega (Sievers and Higgins 2021)version 1.2.4 allowing for five iterations (--iterations 5). Gaps in the resulting alignment were trimmed using trimAl v1.4.rev15 and the -gappyout option. The phylogenetic relationship among proteins was inferred from the trimmed alignment using FastTree (Price et al. 2009) under the Whelan-And-Goldman 2001 model after 1,000 bootstraps (-boot 1000 and -wag options). Plots were performed in R v 4.1.2 using ggtree v3.9.1.

### Estimation of retrotranposon insertion time and nucleotide divergence time

To get an estimation of the insertion date of the transposable elements present in the MAT-proximal non-recombining region, we applied two complementary methods. We first used the divergence between the LTR sequences in retrotransposons, as these LTR sequences at their edges are identical at the time of TE insertion and then diverge with time. For this, we used a *de-novo* prediction of LTR retrotransposons using LTRharvest GenomeTools 1.6.2 (Ellinghaus et al. 2008; Gremme et al. 2013) and LTR_Finder v1.07 (Xu and Wang 2007). To prepare data for LTRharvest, we first created enhanced suffix for the M1400 and M6697 genome assemblies using the GenomeTools Suffixerator (-dna -suf -lcp -bwt -bck -mirrored parameters). We ran LTRharvest using two pipelines, designed to identify LTR retrotransposons with a minimum and maximum LTR length of 100 bp and 7000 bp respectively and at least 85% identity between two LTR regions, with and without canonical TGCA motifs, respectively: i) -minlenltr 100 -maxlenltr 7000 -mintsd 4 -maxtsd 6 -similar 85 -vic 10 -seed 20 -motif TGCA -motifmis 1; ii) -minlentltr 100, -maxlenltr 7000, -mintsd 4, - maxtsd 6, -similar 85, -vic 10, -seed 20. Similarly, we ran LTR_Finder on the M1400 and M6697 genome assemblies to retrotransposons with both TGCA and non-TGCA motifs and a minimum and maximum LTR length of 100 bp and 7000 bp respectively and at least 85% identity between two LTR regions (-D 15000, -d 1000, -l 100, -L 7000, -p 20, -C, -M 0.85). Finally, we used LTR_retriever v2.8.5 (Ou and Jiang 2018) with default parameters to filter out false positive LTR candidates identified by LTRharvest and LTR_Finder and get an estimation of each element insertion date.

As a second method to estimate the insertion date of the transposable elements present in the MAT-proximal non-recombining region, we used the set of curated consensus sequences from (Demené et al. 2022) to annotate the inversion sequence or its surroundings. We first used samtools faidx v1.9 (Li et al. 2009) to extract the sequence corresponding to inversion and their 50 kb surroundings in M1400 and M6697 genome assemblies. We annotated the sequences corresponding to the inversions and the concatenated 50 kb surroundings in both isolates using RepeatMasker version 4.1.5 and rmblast as search engine (v2.10.0) with -no_is -pa 20 -cutoff 250 -a parameters. Finally, we parsed the RepeatMasker .out file using the helped script parseRM_merge_interrupted.pl and omitting Simple_repeat and Low_complexity regions (https://github.com/4ureliek/Parsing-RepeatMasker-Outputs). We then built a summary of the alignments using the RepeatMasker helper script buildSummary.pl and calculated sequence divergence using the calcDivergenceFromAlign.pl script to finally render results with the createRepeatLandscape.pl from the same helper suite.

### Comparative genomics analyses

Genome synteny between long reads assemblies were studied using the nucmer v3.1 program (Marçais et al. 2018). Outputs were plotted with the R programs ggplot2 (Wickham 2009), genoPlotR (Guy et al. 2010) and RIdeogram (Hao et al. 2020). The dotplot in the putative centromere region of the M1400 mating-type contig was performed using the online megablast alignment tool available at https://blast.ncbi.nlm.nih.gov/(last accessed 13^th^ May 2024).

To study gene disruption, synonymous and non-synonymous divergence (d_S_ and d_N_) and testing introgression in the MAT-proximal region, we build orthology relationships for genes of the genome assemblies of *C. parasitica* strains M1400, M6697, XIM9508, MRC10, DUM-005, ESM15 and EP155 and included three genomes of the closely related species *Cryphonectria japonica* (IF-6), *Cryphonectria carpinicola* (CS3), and *Cryphonectria naterciae* (M3664). Genome, gene annotation and species tree of these closely related species were previously published by (Stauber et al. 2020). Genome data were retrieved from NCBI bioproject number PRJNA644891 and accession IDs JACWRX000000000 for IF-6, JACWRQ000000000 for CS3 and JACWST000000000 for M3664. We run OrthoFinder v2.3.7 (Emms and Kelly 2019) analysis on protein sequences. We used the translatorX v1.1 program (Abascal et al. 2010) with default parameters that use a codon-based approach to align orthologous gene coding sequences. To compute d_S_ and d_N_ vaues of one-to-one orthologs between pairs of genome assemblies, we use the yn00 program of PAML (Yang 2007). Estimation of divergence time between haplotypes was performed using computed gene d_S_ values and the formula Tgenerations = dS/2μ. We used previous estimates of substitution rates in fungi (Kasuga et al. 2002; Taylor and Berbee 2006) and considered that *C. parasitica* undergoes one generation a year (Guerin et al. 2001). We build gene coding sequences trees with the outgroup genomes in the MAT-proximal region using iqtree2 v2.2.2.6 (Minh et al. 2020) with 1000 bootstraps and used the Newick Utilities (https://github.com/tjunier/newick_utils) for displaying phylogenetics tree.

## Acknowledgments

This work was supported by the European Research Council (ERC) EvolSexChrom (832352) grant and a Fondation Louis D grant from the Institut de France to TG, and by the ANR PIA grant # ANR-20-IDEES-0002 to F.E.H. We acknowledge the GenOuest bioinformatics core facility (https://www.genouest.org) for providing the computing infrastructure, (GPU) for genome annotation". We thank Jeanne Ropars, Aaron Vogan and Marie Foulongne for discussions and suggestions on the analyses. The authors declare that they have no competing interests.

TG and FEH conceptualized the study and acquired funding. TG supervised the study. AL, AS, CD, AD, SP, DC, LS and TB contributed to data and their availability. AL and AS performed *in vitro* culture and DNA extraction. FEH, RRDLV, QR, JPV, LS and TB analyzed genomes. FEH and TG wrote the original draft. All authors edited the manuscript.

The raw data and the new assemblies produced in this study will be published pending scientific review.

