## Supplementary material for "An old inversion polymorphism involving giant mobile elements in an invasive fungal pathogen"

### Supplementary Tables

**Table S1: List of the *Cryphonectria parasitica* strains sequenced previously with Illumina used in this study.** Information on the strain name, country of sampling, mating type, and genetic cluster was retrieved from (Demené et al. 2019; Stauber et al. 2021; Stauber et al. 2022). Information on the haplotype in the MAT-proximal region (MAT-Prox1 or MAT-Prox2) was inferred from allele frequencies analyses in the MAT-proximal region.

**Table S2: Statistics of SNPs (single nucleotide polymorphisms) in high linkage disequilibrium (LD; $r^2$ >0.9) per scaffold in the *Cryphonectria parasitica* CL1 genetic cluster and the Swiss 1990 and the Swiss 2019 populations:** median and maximal distance between SNPs with r2>0.9 (in bp), number and percentage of SNP pairs with $r^2$ >0.9, and number of SNP pairs.

**Table S3: Statistics of the produced raw reads and the obtained assemblies for the M1400 and M6697 genomes using PacBio sequencing and the XIM9508, MRC010 and DUM005 genomes using MINION sequencing.** Number of contigs, size of the largest contig (in bp), total assembly size (in bp), percentage of GC, N50, N90, Busco completion, percentage of the assemblies masked by repeats, number of raw reads, median read length, read length N50 and total bases produced are indicated. Statistics of the previously published assemblies EP155 (Crouch et al. 2020) and ESM15 (Demené et al. 2022) are indicated.

**Table S4: Diversity and divergence statistics in the MAT-proximal region and other regions in the *Cryphonectria parasitica* 1990 Swiss population.** All statistics were computed per 50-kb window overlapping over 10 kbp for strains of the MAT-Prox1 haplotype and strains of the MAT-Prox2 haplotype in the 1990 Swiss population using the M1400 genome as reference. **A.** Median values of nucleotide diversity within pools of strains of each MAT-proximal haplotype, relative divergence ($F_{ST}$) between strains of the two MAT-proximal haplotypes, and Tajima's D within the pool of all strains; Windows containing fewer than 5 SNPs were removed from the analysis. **B-C.** Median and mean values of $d_S$ (B) and $d_N/d_S$ (C) between M1400 and M6697 genomes. Number of gene pairs with computed values are indicated. **D.** Estimates of divergence time between MAT-proximal haplotypes obtained for different substitution rate estimates and using either the mean $d_S$ values between the MAT-proximal haplotypes.

**Table S5: Summary of the analysis of variance (ANOVAs) of transposable element (TE) load (TE occupation in bp) depending on TE family, genomic region, strain and interactions between factors when significant.** The model of the ANOVA was TE load ~ TE family + genomic region + strain + genomic region : TE family.

**Table S6: Expression and functional annotations for genes located in the MAT-proximal region of the EP155 *Cryphonectria parasitica* strain (MAT-Prox1).** The annotation of the EP155 genome found on the JGI website (https://genome.jgi.doe.gov/portal/) was used. Columns give the start and end positions, as well as strand of the predicted genes, on the scaffold_2 (EP155 genome), the presence of secretion signal (SignalP), the presence of transmembrane helices (predicted by TMHMM) the protein family (Pfam), the InterproScan domain (IPR_description), and the gene ontology (GO) of the predicted encoded protein. The three last columns indicate genes that have support from RNAseq data (Chun et al. 2020), genes that were found differentially expressed in the studies by (Chun et al. 2020) and by (Belov et al. 2021). Genes specific to M6697 haplotypes are indicated in grey and gene with DUF3435 domain (putative Starship captain) are shown in red.

**Table S7: Functional annotations for genes located in the MAT-proximal region of the M6697 *Cryphonectria parasitica* strain (MAT-Prox1).** The annotation of the M6697 genome was performed in this study. Columns give the gene id, the orthologous group id, start and end positions, as well as strand of the predicted genes, on the tig00000060 (M6697 genome), the the protein family (Pfam), the InterproScan domain (IPR_description), and the ENOG domain of the predicted encoded protein. Genes specific to M6697 haplotypes are indicated in grey and gene with DUF3435 domain (putative Starship captain) are shown in red.

**Table S8: Functional annotations for genes located in the MAT-proximal region.** The annotation of the EP155 genome found on the JGI website (https://genome.jgi.doe.gov/portal/) was used. Columns give the start and end positions, as well as strand of the predicted genes, on the scaffold_2 (EP155 genome), the presence of secretion signal (SignalP), the presence of transmembrane helices (predicted by TMHMM) the protein family (Pfam), the InterproScan domain (IPR_description), and the gene ontology (GO) of the predicted encoded protein.

**Table S9: Functional annotations for ortholous genes in the MAT-proximal region of M1400 and M6697 with dN/dS ratio > 1.**

**Supplementary Figures**

**S Figure 1: Linkage disequilibrium (LD) along the contig carrying the mating-type locus in the invasive *Cryphonectria parasitica* Swiss 1990 (A) and Swiss 2019 (B) populations.** LD heatmaps using single nucleotide polymorphisms (SNPs) located on the chromosome carrying the mating-type locus (scaffold_2 of the EP155 genome). The mating-type locus location is shown with a green triangle and the MAT-proximal region lacking recombination is shown with a red arrow. The two high-LD blocks within the MAT-proximal region are shown with orange arrows. SNPs at the limit of the MAT-proximal region and the two high LD blocks were manually highlighted with red dotted lines and an orange line.

**S Figure 2: Linkage disequilibrium (LD) along other scaffolds of the EP155 genome in the invasive *Cryphonectria parasitica* CL1 genetic cluster. A.** Distribution of the SNPs density (per 100 kb) along the scaffold_2 of EP155 genome. SNP density was computed per 100-kb non-overlapping windows. The MAT-proximal region defined from LD analyses is indicated with red arrow and dotted lines. The mating-type locus location is shown with a green triangle and a green dotted line. **B.** LD along the contig carrying the mating-type locus (scaffold_2) for SNPs that are distant of at least 50 kb. The mating-type locus location is shown with a green triangle and the MAT-proximal region lacking recombination is shown with a red arrow. **C.** LD along the scaffold_6. The high-LD values between pairs of distant SNPs on scaffold_6 (blocks at 3-20 kb and 2.151-2.278 kb) are indicated with an empty red square. **D.** Distribution of the distance (bp) between pairs of SNPs with high linkage disequilibrium (LD, i.e. $r^2$ >0.9) along all contigs. The y-axis was log transformed.

**S Figure 3: Genetic structure using single nucleotide polymorphisms (SNPs) in the MAT-proximal region lacking recombination and in other genomic regions in the invasive *Cryphonectria parasitica* Swiss 1990 and 2019 populations. A and C.** Principal component analysis (PCA). Two principal components are presented. Percentage of variance explained by each PC is indicated into brackets. Strains are colored according to their mating type (*MAT1-1* or *MAT1-2*). **B and D.** Neighbor-net tree from a SplitsTree analysis. These analyses were performed in the Swiss 1990 and 2019 populations (European invasive populations introduced from North America) based on: (1) SNPs located within the MAT-proximal region. (2) SNPs located in other regions (i.e. other contigs than the contig carrying the MAT-proximal region). The two clusters corresponding to the MAT-Prox1 et MAT-Prox2 haplotypes are shown with red circles. The MAT-Proximal haplotype of each strain is indicated in Table S1. The number of strains in the MAT-Prox1 cluster is indicated by the letter n.

**S Figure 4: Linkage disequilibrium (LD) analysis in the invasive *Cryphonectria parasitica* Swiss 1990 populations around the mating-type locus in the M1400 newly sequenced genome.** LD heatmaps using single nucleotide polymorphisms (SNPs) located on the chromosome carrying the mating-type locus (tig_000001 of the M1400 genome, only positions farther than 7 Mb are shown). The MAT-proximal region lacking recombination corresponds to the red triangle (pairs of SNPs with high LD, i.e. $r^2$ >0.9). The MAT-proximal region defined from LD analyses and the inversion between M1400 and M6697 genomes are indicated with red and blue arrows, respectively. The mating-type locus location is shown with a green triangle. The two high-LD blocks within the MAT-proximal region are indicated with orange arrows. SNPs at the limit of the MAT-proximal region, the two high-LD blocks and the inversion were manually highlighted with red dotted lines, an orange line and blue dotted lines, respectively.

**S Figure 5: Synteny plot between the M1400 and M6697 genomes (invasive population introduced from North America) on (A) the nine largest contigs of the M1400 genome and (B) the MAT-proximal region and (C) dot plot in the putative centromere region of the M1400 mating-type contig.** A. The contigs carrying the mating-type locus are highlighted with a red rectangle. B. The MAT locus and MAT-proximal region are indicated with a green dotted line and triangle and a red dotted line and arrow, respectively, along the M1400 genome. On panel A and B, blue and red colours indicate the same and opposite strand collinearity, respectively. C. Dot plot in a 1.1 Mb region around the putative centromere region of the M1400 mating-type contig (3.9Mb to 5 Mb of tig_000001 of the M1400 genome).

**S Figure 6: Comparison of diversity and divergence statistics in the MAT-proximal region and other recombining regions of the M1400 genome in the invasive *Cryphonectria parasitica* Swiss 1990 populations.** A. Relative divergence ($F_{ST}$) between strains of the MAT-Prox1 and MAT-Prox2 haplotypes; B. Nucleotide diversity within pools of strains for each

MAT-Proximal haplotype; C. Tajima's D for all strains pooled and within pools of strains for each MAT-Proximal haplotype;

**S Figure 7: Maximum likelihood tree of the 4 putative Starship captains of the M1400 Cryphonectria parasitica strain and 1222 representative of Starship captains classified in 11 families** (Gluck-Thaler and Vogan 2024). The putative captain M1400bis_01g001231.1|967 is the one clustering with the Arwing family ; the three other captains cluster with the Phoenix family.

**S Figure 8: Comparison of synonymous divergence ($d_S$) and non-synonymous divergence (dN) between high quality genomes of *Cryphonectria parasitica* strains from the native and introduced range. A to D.** Boxplots of the distribution of dS values (left) and $d_N/d_S$ ratio values (right) in the MAT-proximal region and other recombining regions for four pairwise genome comparisons: **A.** M6697 vs M1400 (European invasive strains originating from North America); **B.** MRC10 (European invasive strain originating directly from Asia) vs M1400; **C.** MRC10 vs M6697; **D.** ESM15 vs XIM9508 (native range); **E**. Distribution of dS values along the M1400 mating-type contig for the four pairwise genome comparisons. Only dS values of genes pairs having orthologous gene to M1400 genes and dS values <1 are shown. The mating-type locus and the MAT-proximal region are indicated with a green and red dotted lines.

**S Figure 9: Synteny plots, gene densities and transposable element content of the MAT-proximal region of the high-quality genome assemblies M1400 or M6697 (Switzerland, introduced range, invasion from US) with XIM9508 and ESM15 (Asia, native range), MRC10 (Europe, introduced range, invasion directly from Asia), DUM-005 and EP155 (North America, introduced range). A.** Synteny and gene density of XIM9508 and M1400 mating-type contigs. Blue links show colinear regions in the MAT-proximal region. Grey links show other regions of the mating-type contigs. The mating-type locus is located with a green diamond. Gene density tracks is shown with a color gradient (blue with low density, orange with high density). **B.** Dotplot of the MAT-proximal region of M1400 versus XIM9508. **C.** Dotplot of the MAT-proximal region of M1400 versus ESM15. **D.** Dotplot of the MAT-proximal region of M1400 versus MRC10. **E.** Dotplot of the MAT-proximal region of M1400 versus DUM-005. **F.** Dotplot of the MAT-proximal region of M6697 versus EP155. The MAT-proximal region is indicated with red arrows. The mating-type locus location is shown with a green triangle. Transposons larger than 5 kb are shown with orange rectangles. **G.** Relative percentage of each TE annotation (in %) in the non-recombining MAT-proximal region and other recombining regions of M6697 and M1400 genomes.

**S Figure 10: Neighbor-net tree from SplitsTree analyses in additional invasive and native *Cryphonectria parasitica* populations.** In each panel based on SNPs in the MAT-Proximal region (1) or other contigs than the mating-type chromosome (2). **A.** North American (US) invasive strains (n=8; (Demené et al. 2019); circled in purple) and strains from the Swiss 2019 invasive population, the MAT-Prox1 and MAT-Prox2 haplotype being indicated; Mating-type allele information is indicated only for the North American strains (the mating-type is associated with the MAT-Proximal haplotypes in the Swiss population); Two haplotypes are found among strains from US. **B.** Strains from the S12 European invasive lineage (n=105; (*30*)) and the strains M1400 (MAT-prox2) and M6697 (MAT-prox1) from the CL1 genetic cluster. All strains of the S12 cluster have the MAT-1 mating-type allele and the MAT-Prox1 haplotype. **C.** European clonal lineages, with identification of those originating directly from Asia and those coming from America, (n=21; (Demené et al. 2019)) and strains from the Swiss 2019 population, the MAT-Prox1 and MAT-Prox2 haplotypes being indicated. The mating type information is indicated only for clonemates with introgression at the mating-type locus. Both MAT-Prox1 and MAT-Prox2 haplotypes were found among clonemates of the lineages re092 and re103. D. The six strains with high-quality genome assemblies (M6697, M1400, DUM005 (Swiss and North America, introduced range), MRC010 (Europe, introduced range from a second introduction wave), ESM15 and XIM9508 (Asia, native range)) with multiple strains of the CL1, CL2, CL3, CL4 clusters. The MAT-Prox1 and MAT-Prox2 haplotype detected in CL1 cluster and the two identified genomic rearrangements are indicated.

Fig S1

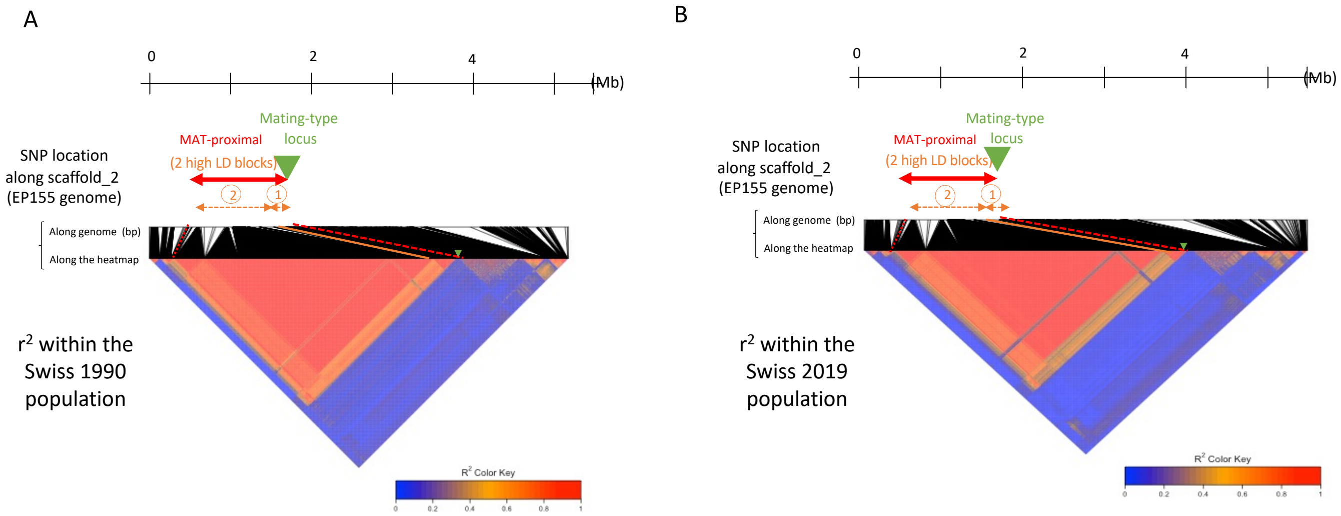
A
0
2
4
(Mb)
Mating-type
locus
MAT-proximal
(2 high LD blocks)
SNP location
along scaffold_2
(EP155 genome)
2
1
Along genome (bp)
Along the heatmap
r² within the
Swiss 1990
population
R² Color Key
0
0.2
0.4
0.6
0.8
1
B
0
2
4
(Mb)
Mating-type
locus
MAT-proximal
(2 high LD blocks)
SNP location
along scaffold_2
(EP155 genome)
2
1
Along genome (bp)
Along the heatmap
r² within the
Swiss 2019
population
R² Color Key
0
0.2
0.4
0.6
0.8
1

Fig S2

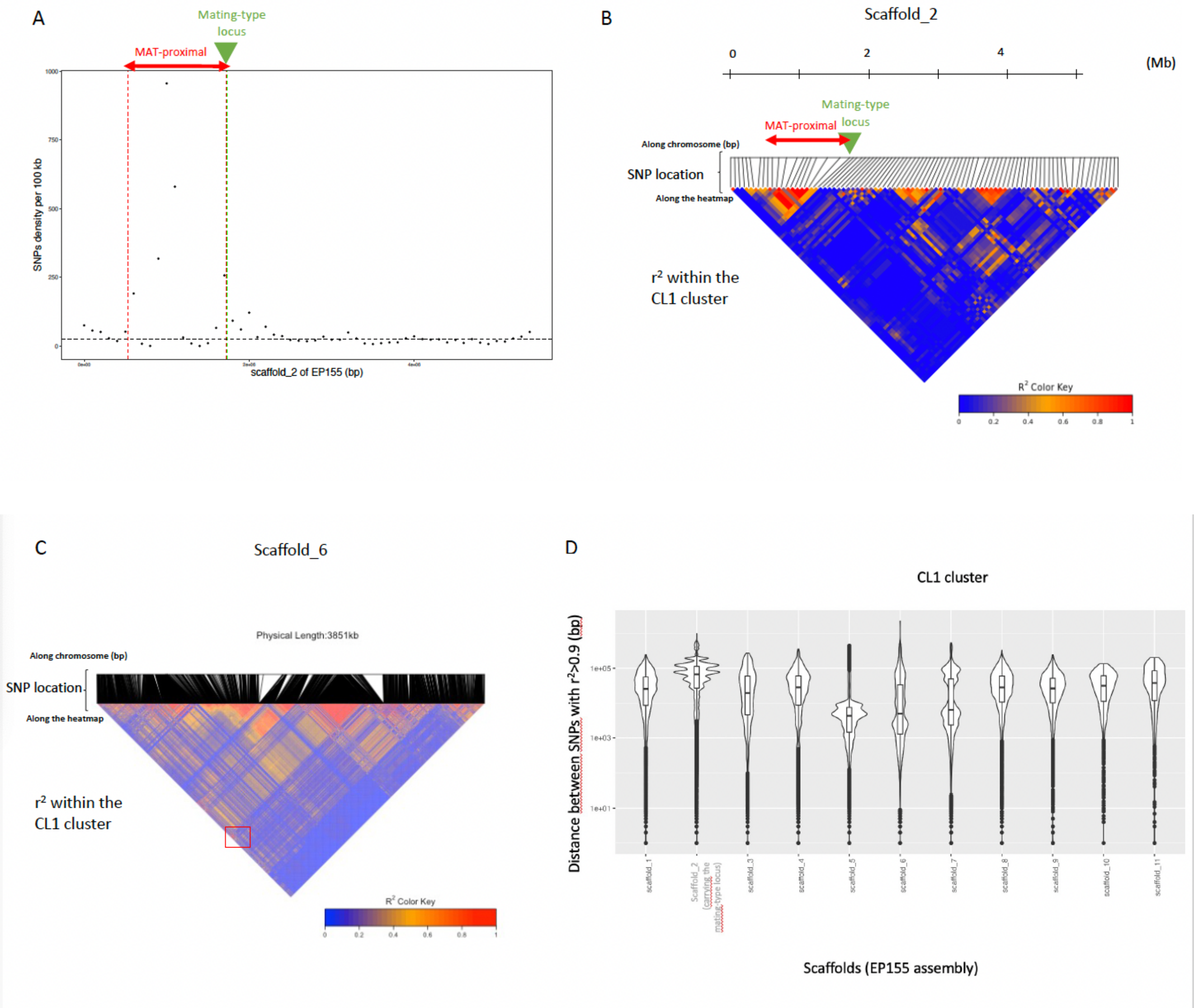
A
Mating-type locus
MAT-proximal
SNPs density per 100 kb
1000
750
500
250
0
scaffold_2 of EP155 (bp)
B
Scaffold_2
0
2
4
(Mb)
Mating-type locus
MAT-proximal
Along chromosome (bp)
SNP location
Along the heatmap
r² within the CL1 cluster
R² Color Key
0
0.2
0.4
0.6
0.8
1
C
Scaffold_6
Physical Length:3851kb
Along chromosome (bp)
SNP location
Along the heatmap
r² within the CL1 cluster
R² Color Key
0
0.2
0.4
0.6
0.8
1
D
CL1 cluster
Distance between SNPs with r²>0.9 (bp)
1e+05
1e+03
1e+01
scaffold_1
Scaffold_2 (carrying the mating-type locus)
scaffold_3
scaffold_4
scaffold_5
scaffold_6
scaffold_7
scaffold_8
scaffold_9
scaffold_10
scaffold_11
Scaffolds (EP155 assembly)

Fig S3

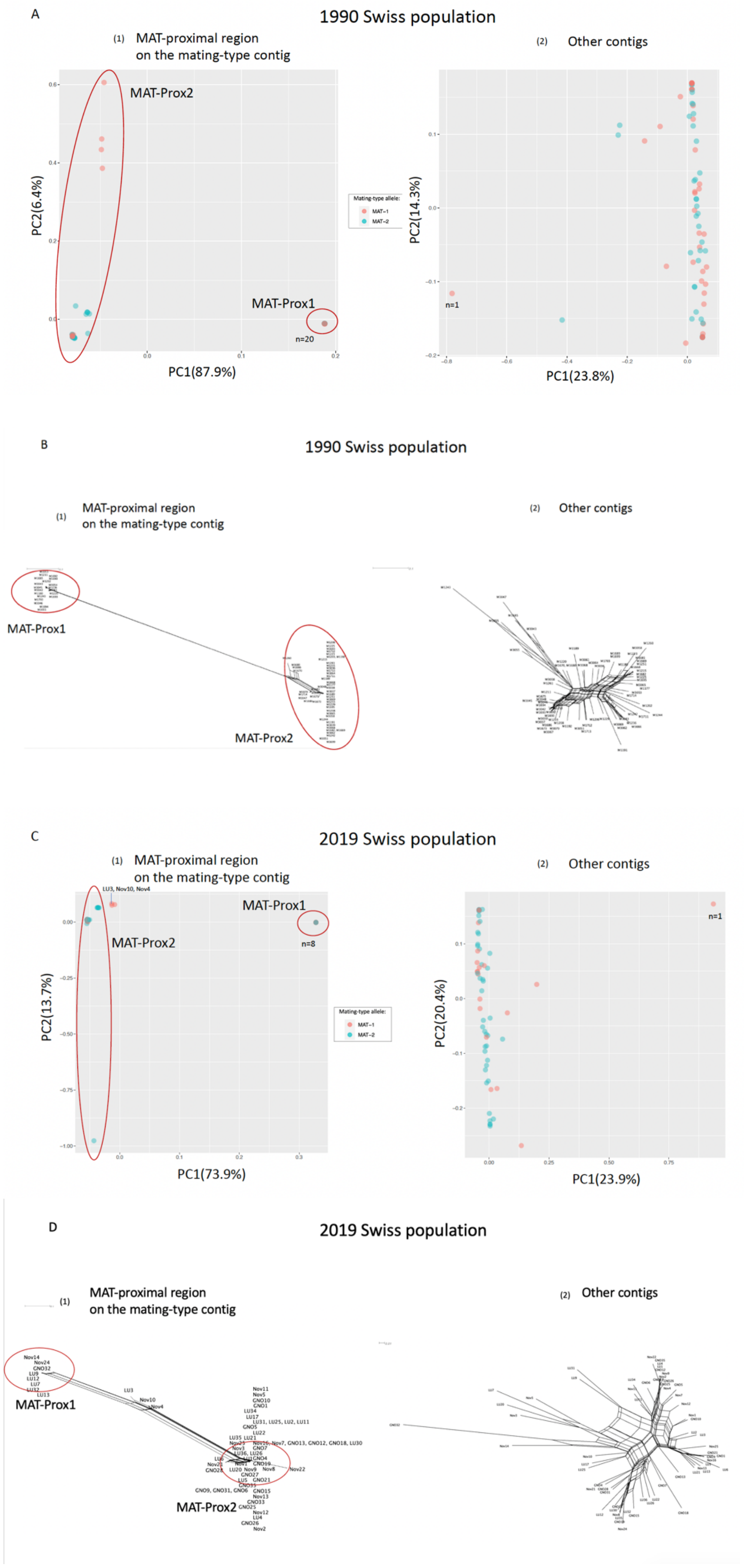
A
1990 Swiss population
(1) MAT-proximal region
on the mating-type contig
(2) Other contigs
MAT-Prox2
MAT-Prox1
n=20
PC1(87.9%)
PC2(6.4%)
Mating-type allele:
MAT-1
MAT-2
n=1
PC1(23.8%)
PC2(14.3%)
B
1990 Swiss population
(1) MAT-proximal region
on the mating-type contig
(2) Other contigs
MAT-Prox1
MAT-Prox2
C
2019 Swiss population
(1) MAT-proximal region
on the mating-type contig
(2) Other contigs
LU3, Nov10, Nov4
MAT-Prox1
MAT-Prox2
n=8
PC1(73.9%)
PC2(13.7%)
Mating-type allele:
MAT-1
MAT-2
n=1
PC1(23.9%)
PC2(20.4%)
D
2019 Swiss population
(1) MAT-proximal region
on the mating-type contig
(2) Other contigs
MAT-Prox1
MAT-Prox2

Fig S4

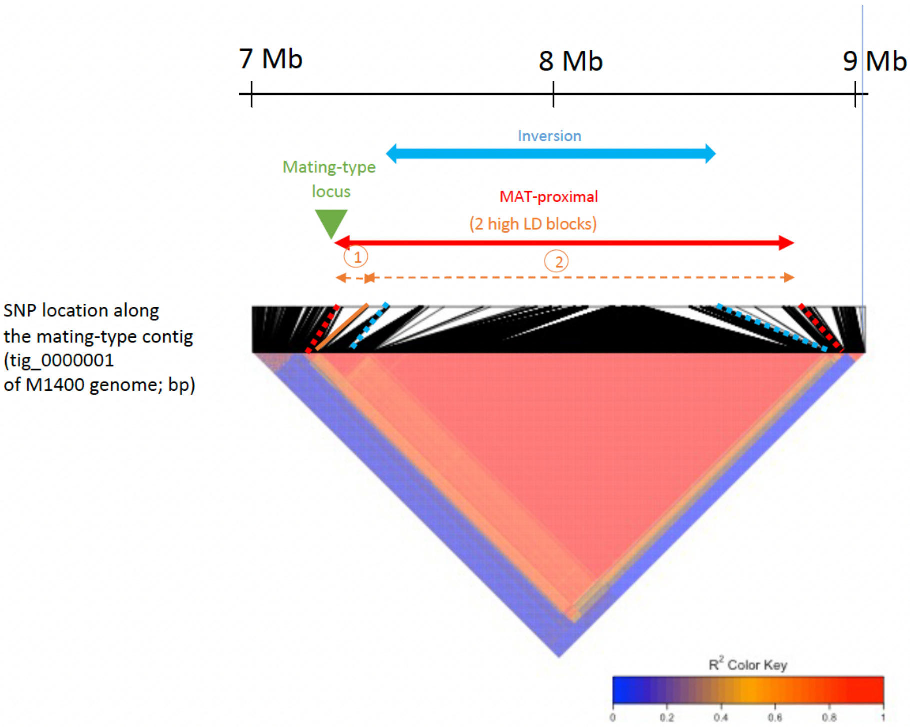
7 Mb
8 Mb
9 Mb
Inversion
Mating-type locus
MAT-proximal
(2 high LD blocks)
1
2
SNP location along the mating-type contig (tig_0000001 of M1400 genome; bp)
R2 Color Key
0
0.2
0.4
0.6
0.8
1

Fig S5

A

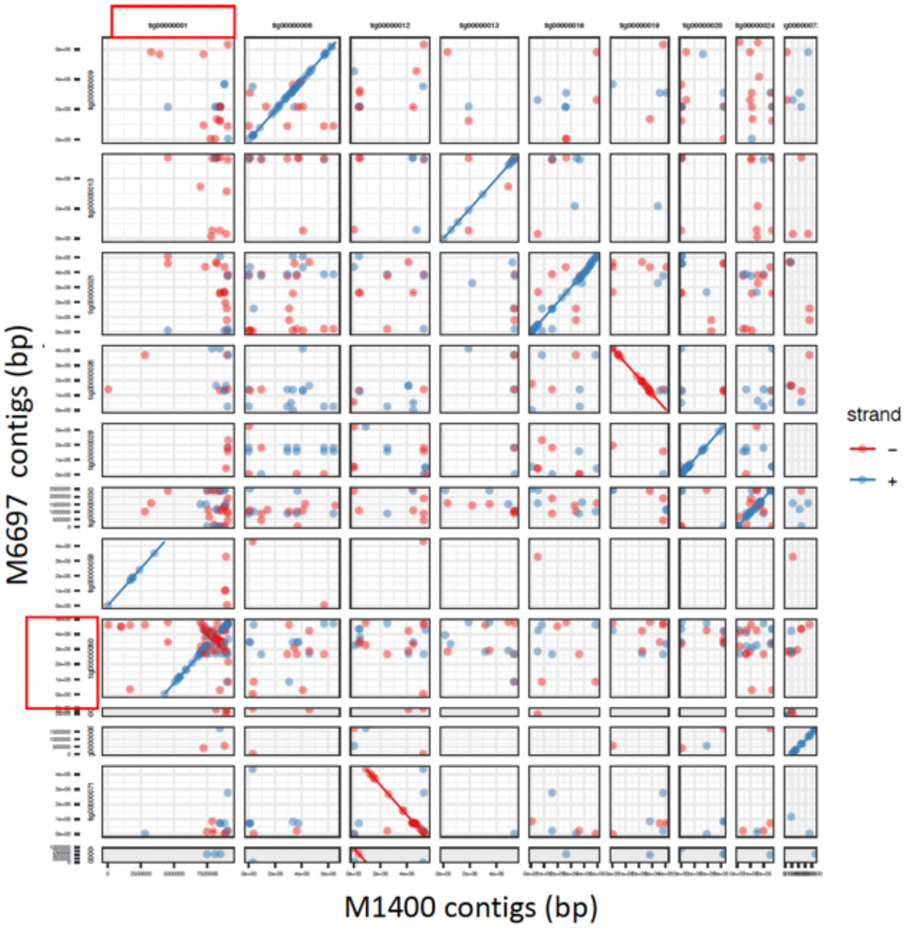

B

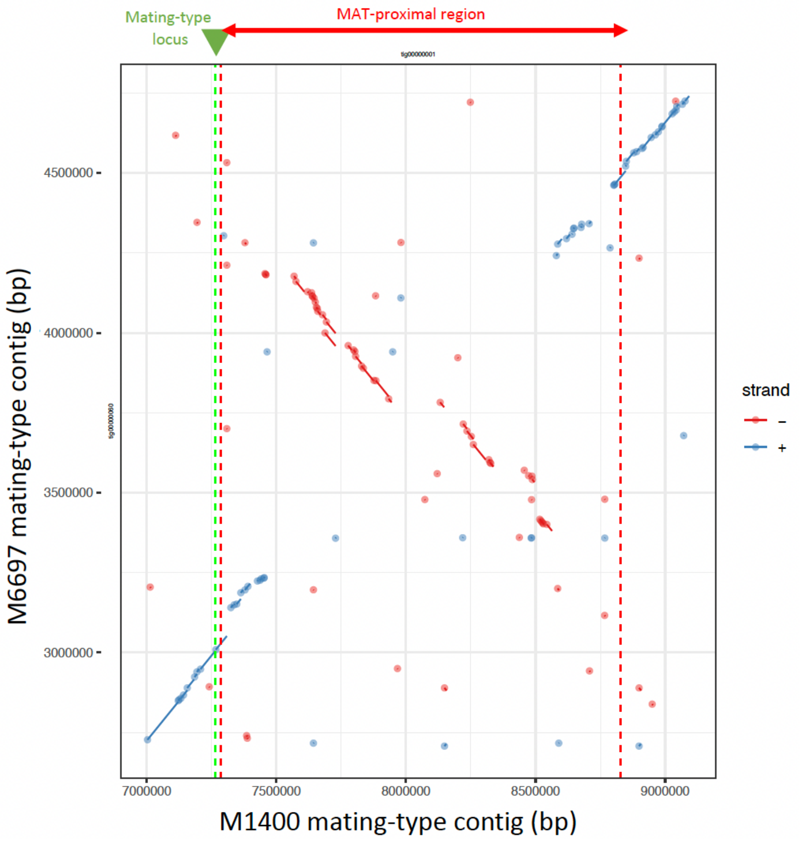
Mating-type
locus
MAT-proximal region
4500000
4000000
3500000
3000000
M6697 mating-type contig (bp)
7000000
7500000
8000000
8500000
9000000
M1400 mating-type contig (bp)
strand
–
+

C

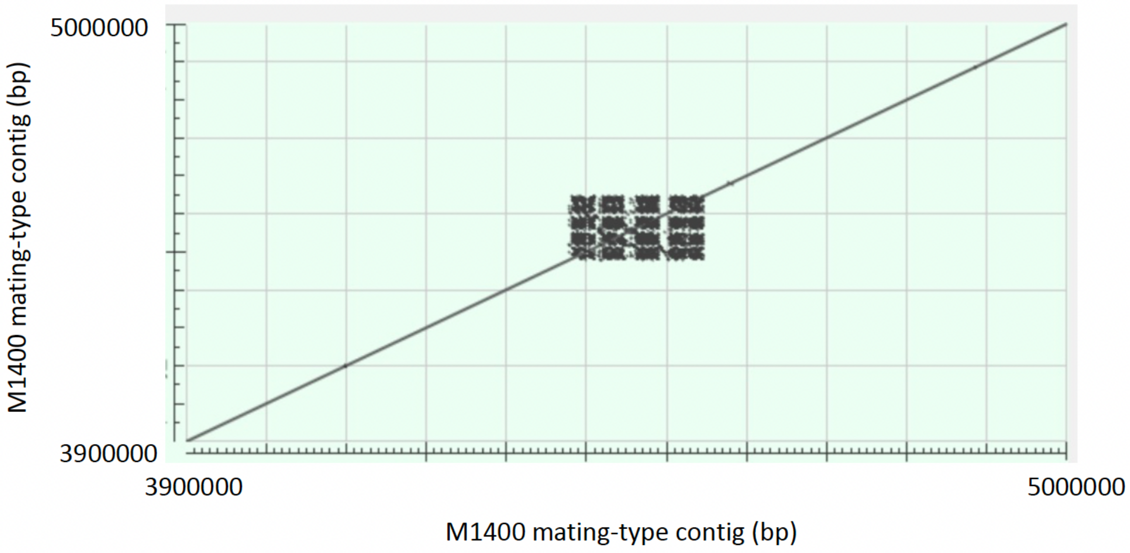
5000000
3900000
M1400 mating-type contig (bp)
3900000
5000000
M1400 mating-type contig (bp)

Fig S6

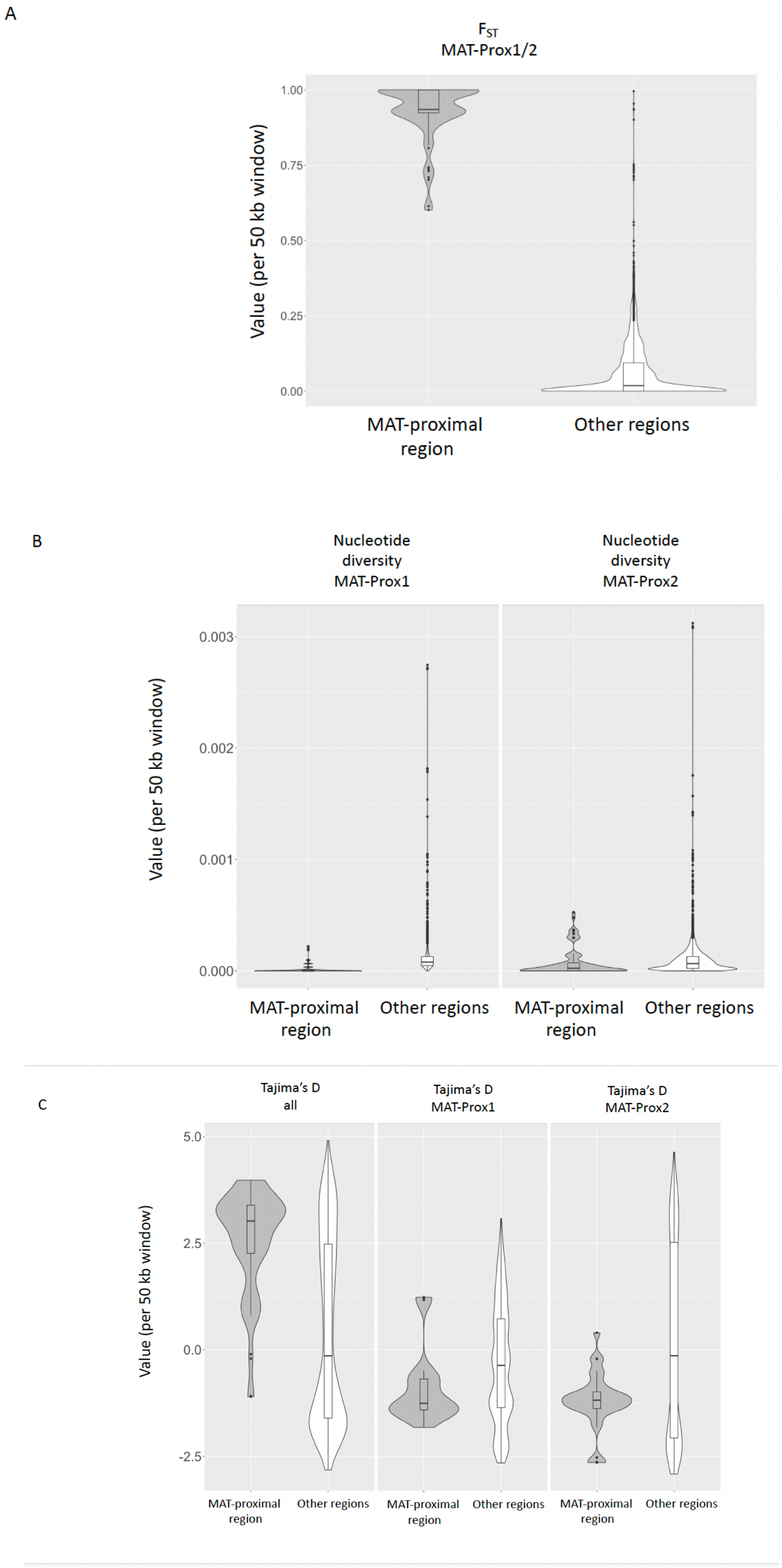
A
$F_{ST}$
MAT-Prox1/2
Value (per 50 kb window)
1.00
0.75
0.50
0.25
0.00
MAT-proximal region
Other regions
B
Nucleotide diversity MAT-Prox1
Nucleotide diversity MAT-Prox2
Value (per 50 kb window)
0.003
0.002
0.001
0.000
MAT-proximal region
Other regions
MAT-proximal region
Other regions
C
Tajima's D all
Tajima's D MAT-Prox1
Tajima's D MAT-Prox2
Value (per 50 kb window)
5.0
2.5
0.0
-2.5
MAT-proximal region
Other regions
MAT-proximal region
Other regions
MAT-proximal region
Other regions

Fig S7

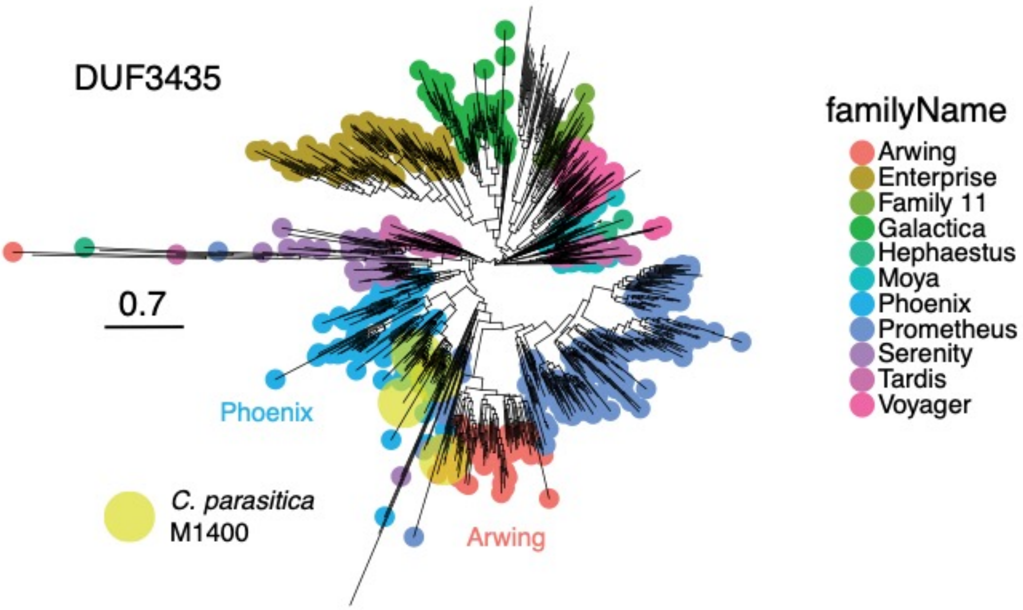
DUF3435
0.7
Phoenix
Arwing
C. parasitica
M1400
familyName
Arwing
Enterprise
Family 11
Galactica
Hephaestus
Moya
Phoenix
Prometheus
Serenity
Tardis
Voyager

Fig S8

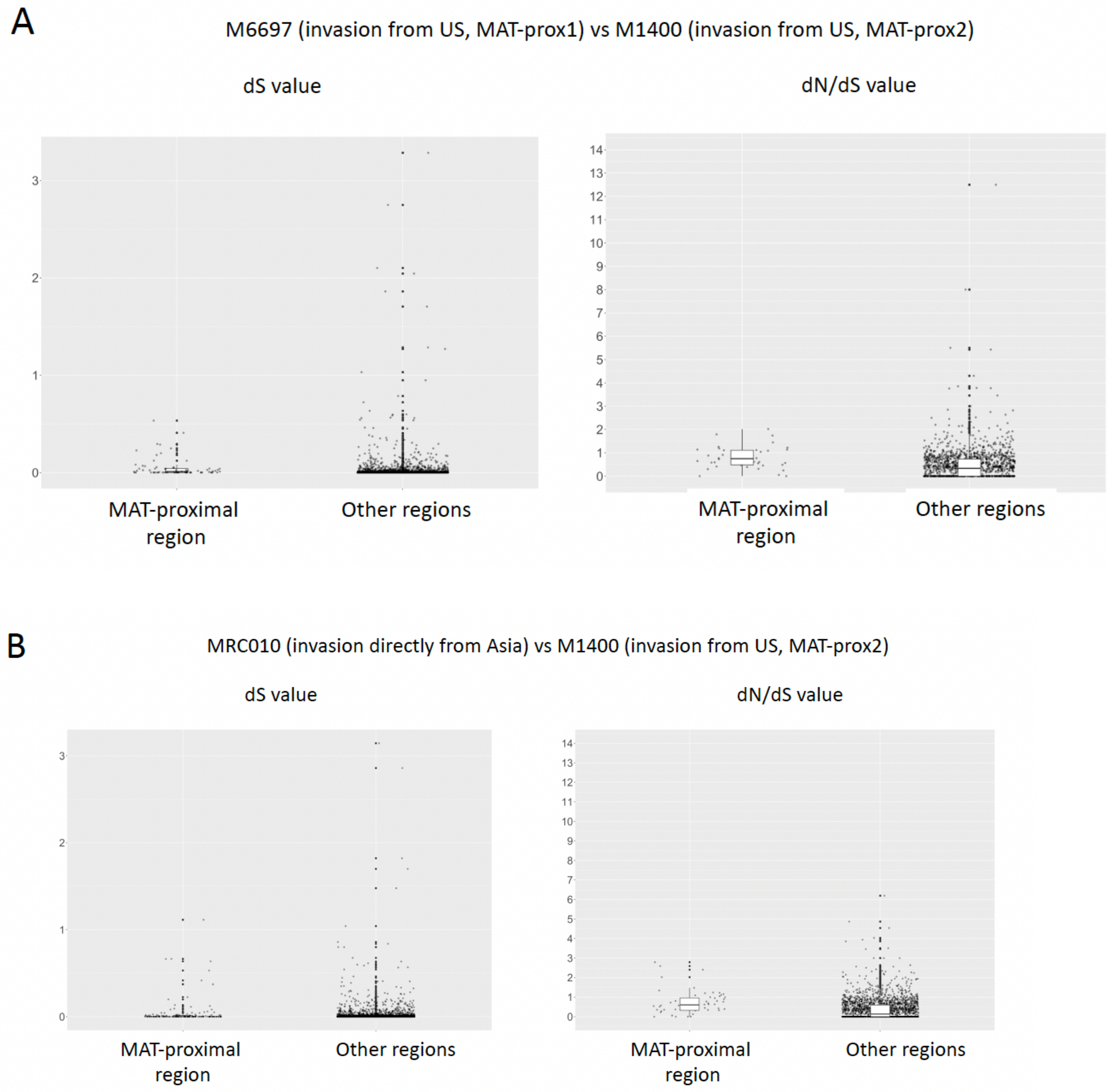
A
M6697 (invasion from US, MAT-prox1) vs M1400 (invasion from US, MAT-prox2)
dS value
dN/dS value
MAT-proximal region
Other regions
MAT-proximal region
Other regions
B
MRC010 (invasion directly from Asia) vs M1400 (invasion from US, MAT-prox2)
dS value
dN/dS value
MAT-proximal region
Other regions
MAT-proximal region
Other regions

C

MRC010 (invasion directly from Asia) vs M6697 (invasion from US, MAT-prox1)

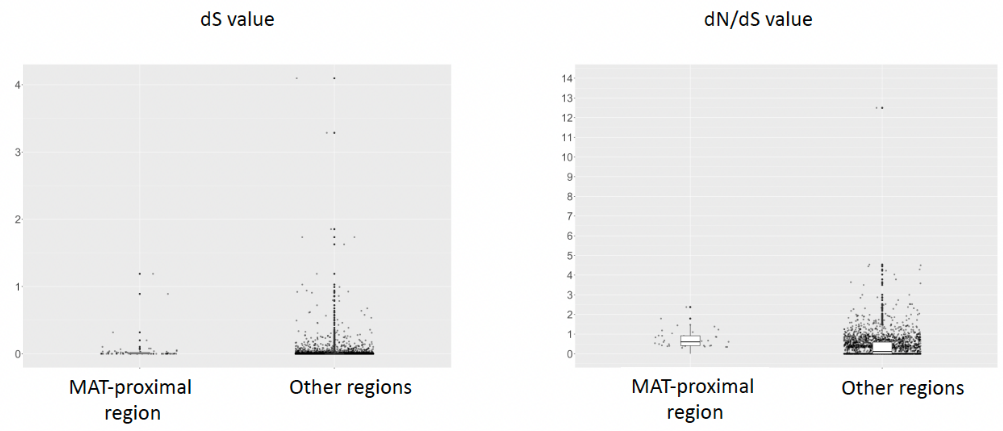


D

ESM15 vs XIM9508 (native range)

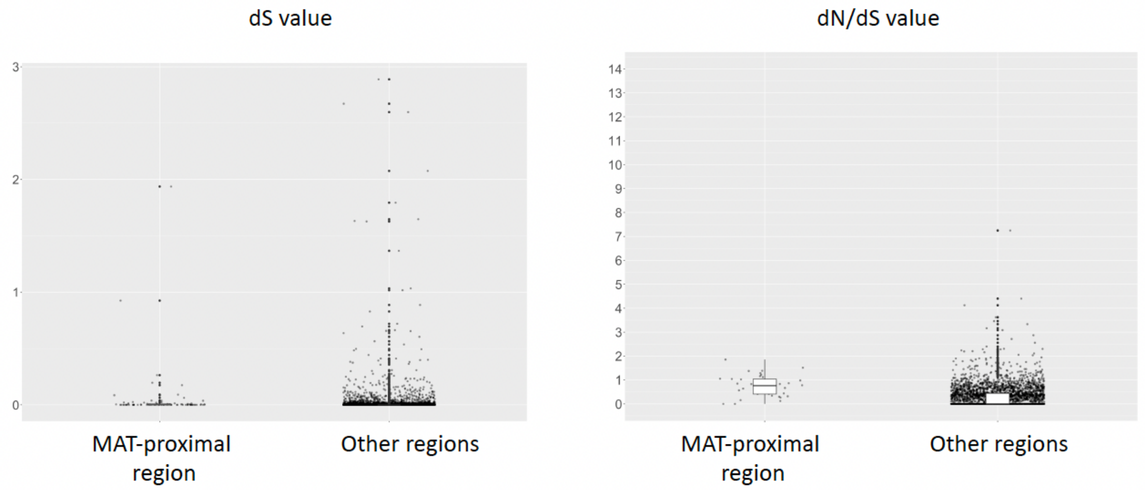

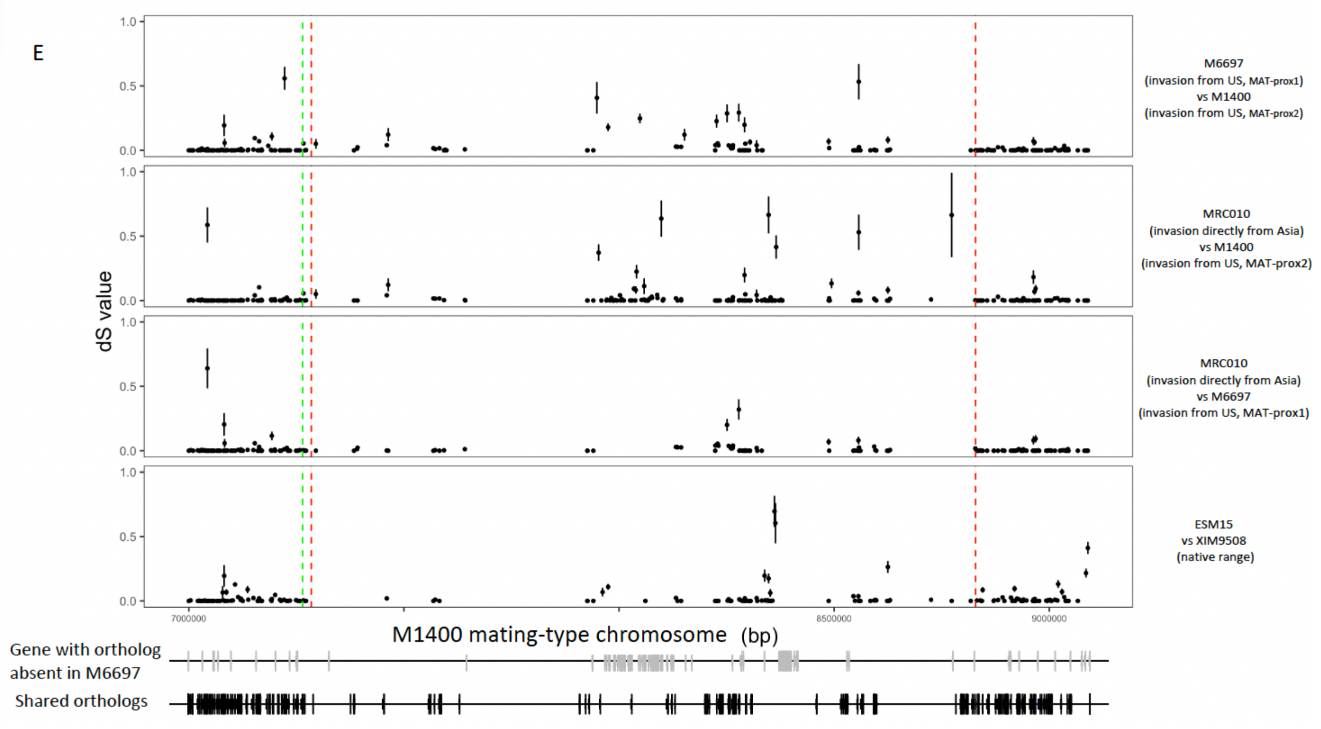
E
dS value
M6697
(invasion from US, MAT-prox1)
vs M1400
(invasion from US, MAT-prox2)
MRC010
(invasion directly from Asia)
vs M1400
(invasion from US, MAT-prox2)
MRC010
(invasion directly from Asia)
vs M6697
(invasion from US, MAT-prox1)
ESM15
vs XIM9508
(native range)
M1400 mating-type chromosome (bp)
Gene with ortholog
absent in M6697
Shared orthologs

Fig S9

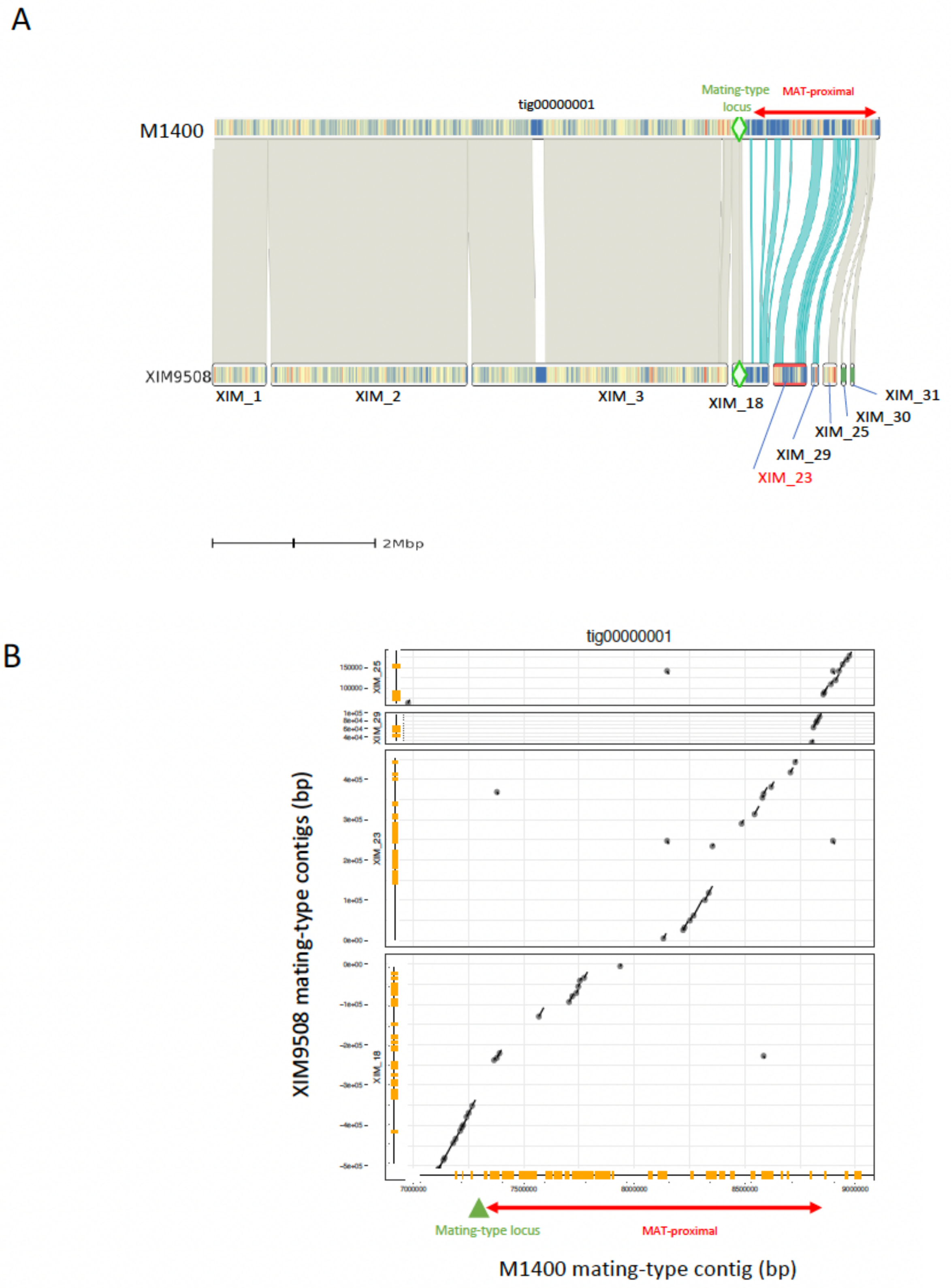
A
M1400
tig00000001
Mating-type locus
MAT-proximal
XIM9508
XIM_1
XIM_2
XIM_3
XIM_18
XIM_31
XIM_30
XIM_25
XIM_29
XIM_23
2Mbp
B
tig00000001
XIM9508 mating-type contigs (bp)
XIM_25
XIM_29
XIM_23
XIM_18
Mating-type locus
MAT-proximal
M1400 mating-type contig (bp)

C

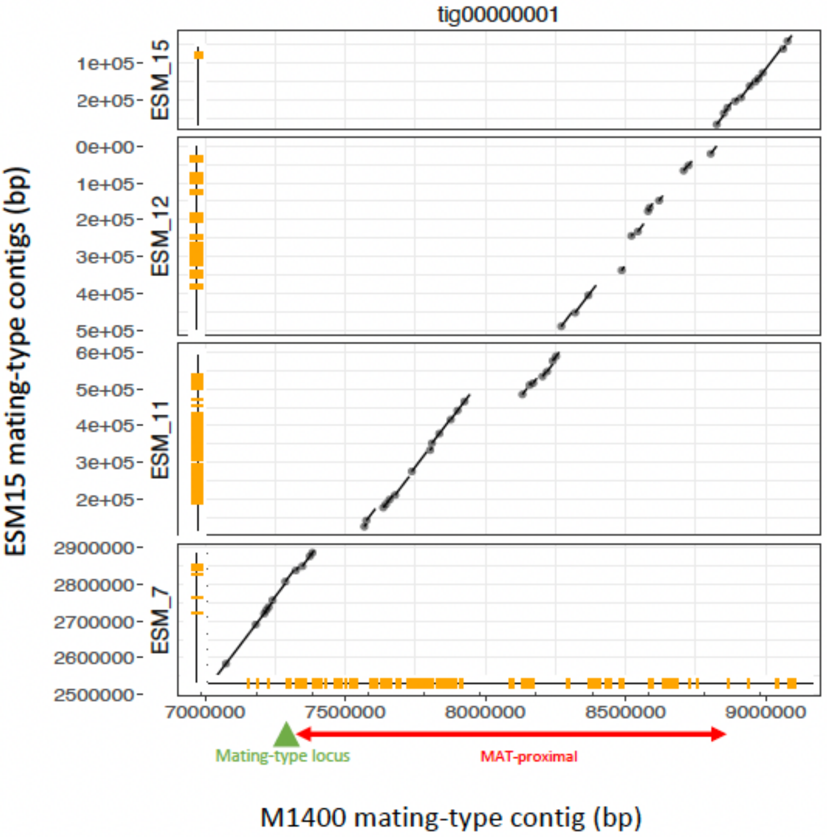
tig00000001
ESM15 mating-type contigs (bp)
ESM_15
1e+05
2e+05
ESM_12
0e+00
1e+05
2e+05
3e+05
4e+05
5e+05
ESM_11
6e+05
5e+05
4e+05
3e+05
2e+05
ESM_7
2900000
2800000
2700000
2600000
2500000
7000000
7500000
8000000
8500000
9000000
Mating-type locus
MAT-proximal
M1400 mating-type contig (bp)

D

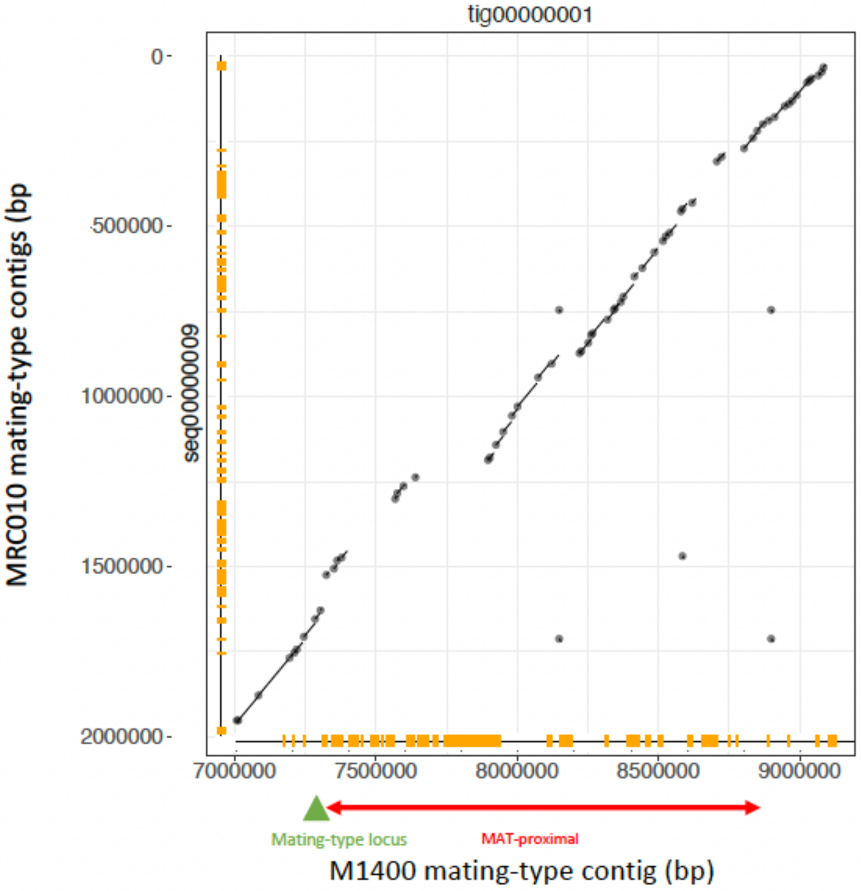
tig00000001
MRC010 mating-type contigs (bp
seq00000009
0
500000
1000000
1500000
2000000
7000000
7500000
8000000
8500000
9000000
Mating-type locus
MAT-proximal
M1400 mating-type contig (bp)

E

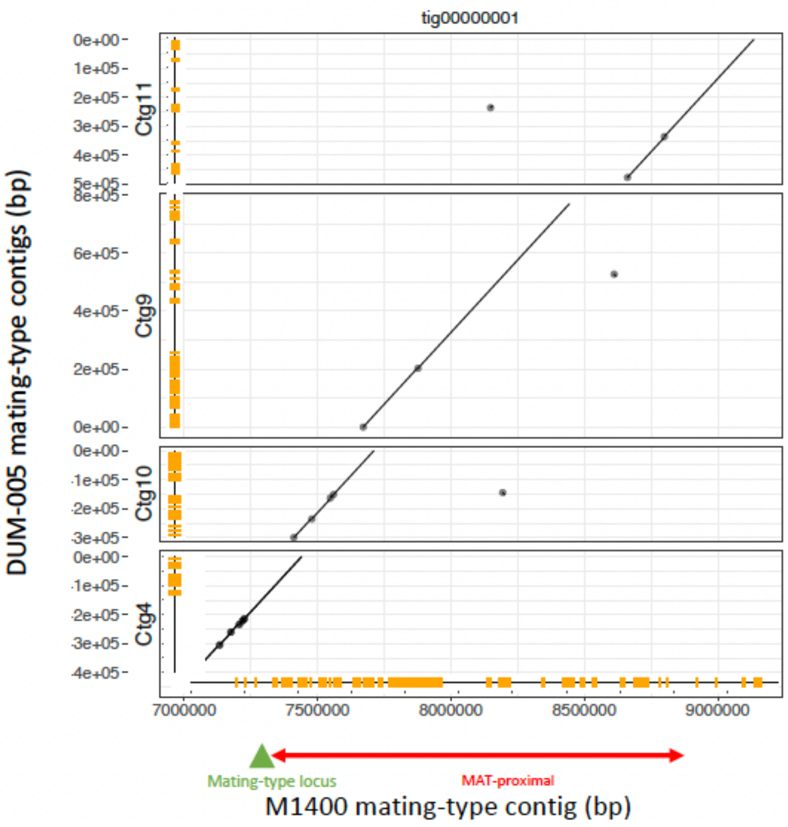
tig00000001
DUM-005 mating-type contigs (bp)
Ctg11
Ctg9
Ctg10
Ctg4
0e+00
1e+05
2e+05
3e+05
4e+05
5e+05
8e+05
6e+05
4e+05
2e+05
0e+00
7000000
7500000
8000000
8500000
9000000
Mating-type locus
MAT-proximal
M1400 mating-type contig (bp)

F

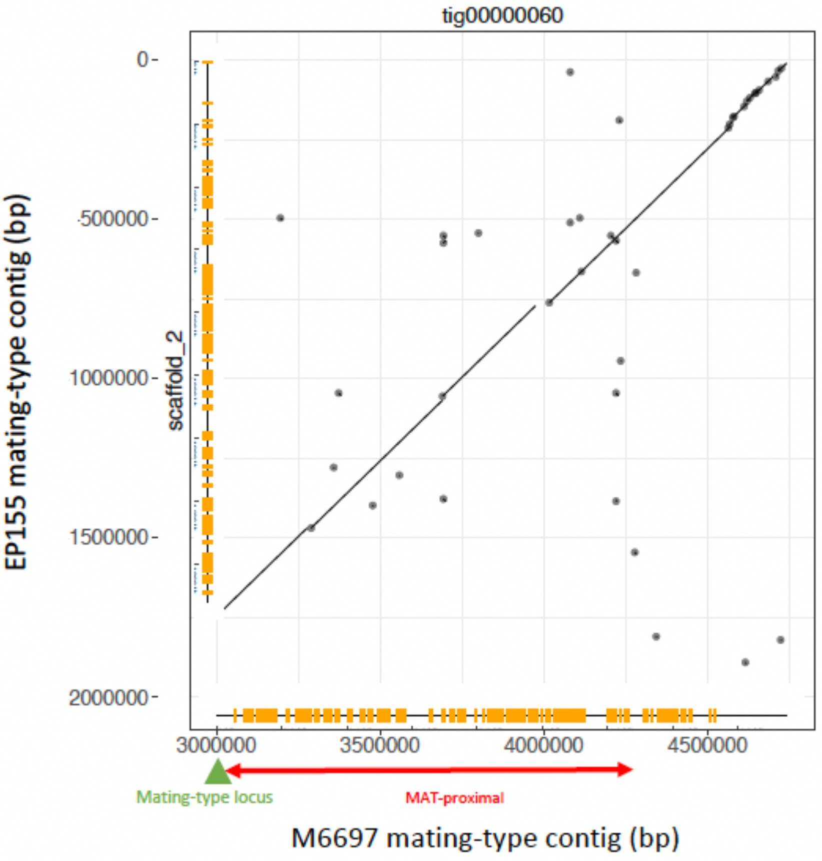
tig00000060
EP155 mating-type contig (bp)
scaffold_2
0
500000
1000000
1500000
2000000
3000000
3500000
4000000
4500000
Mating-type locus
MAT-proximal
M6697 mating-type contig (bp)

G

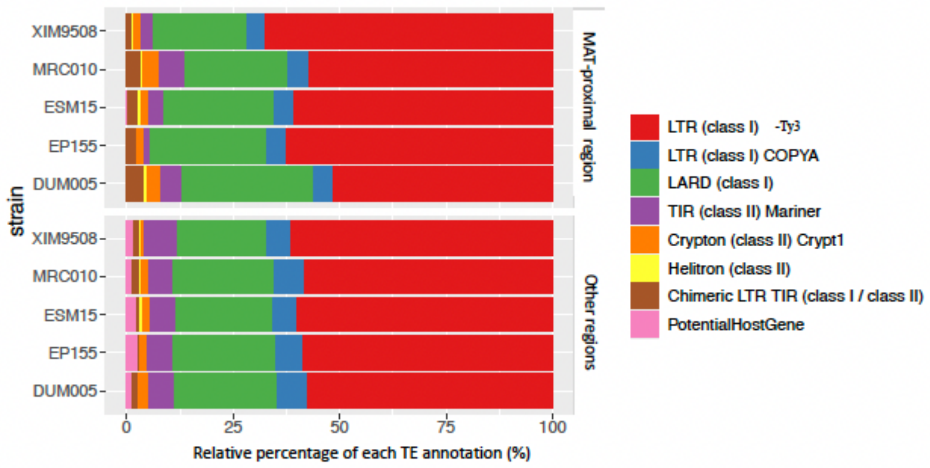
XIM9508
MRC010
ESM15
EP155
DUM005
strain
MAT-proximal region
Other regions
0
25
50
75
100
Relative percentage of each TE annotation (%)
LTR (class I) -Ty3
LTR (class I) COPYA
LARD (class I)
TIR (class II) Mariner
Crypton (class II) Crypt1
Helitron (class II)
Chimeric LTR TIR (class I / class II)
PotentialHostGene

Fig S10

A (1) MAT-proximal region

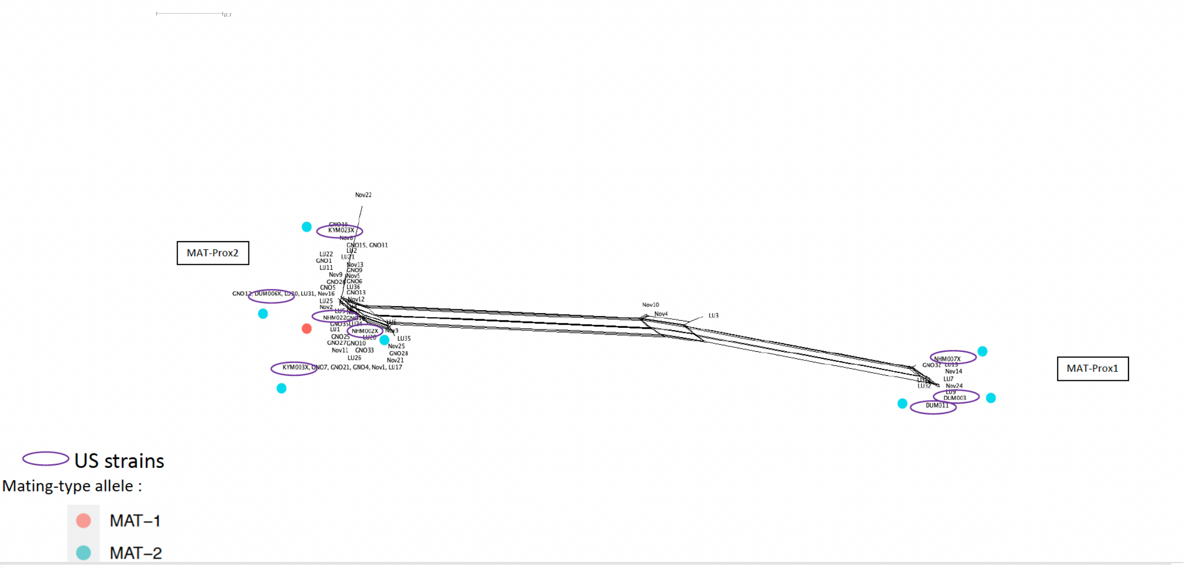


A (2) other contigs

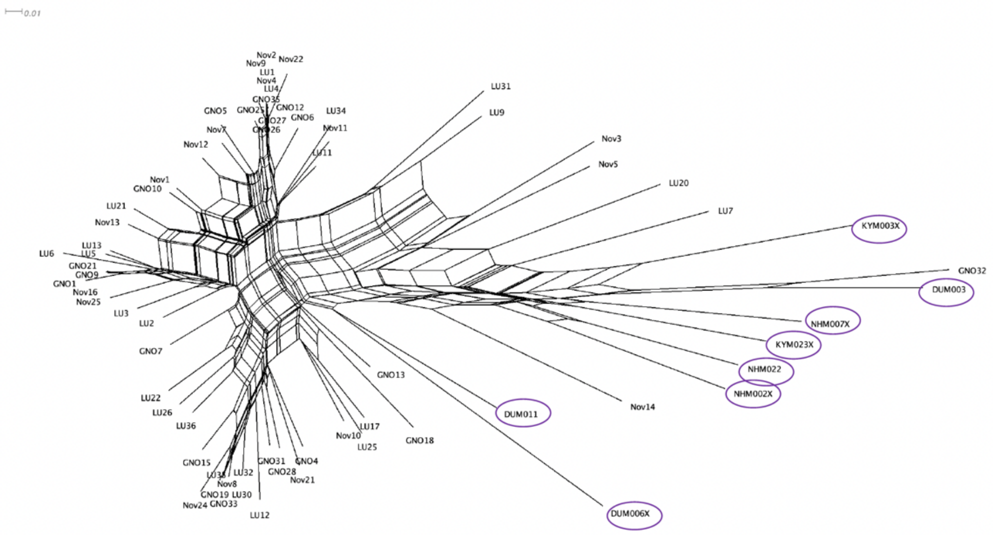

B (1) MAT-proximal region

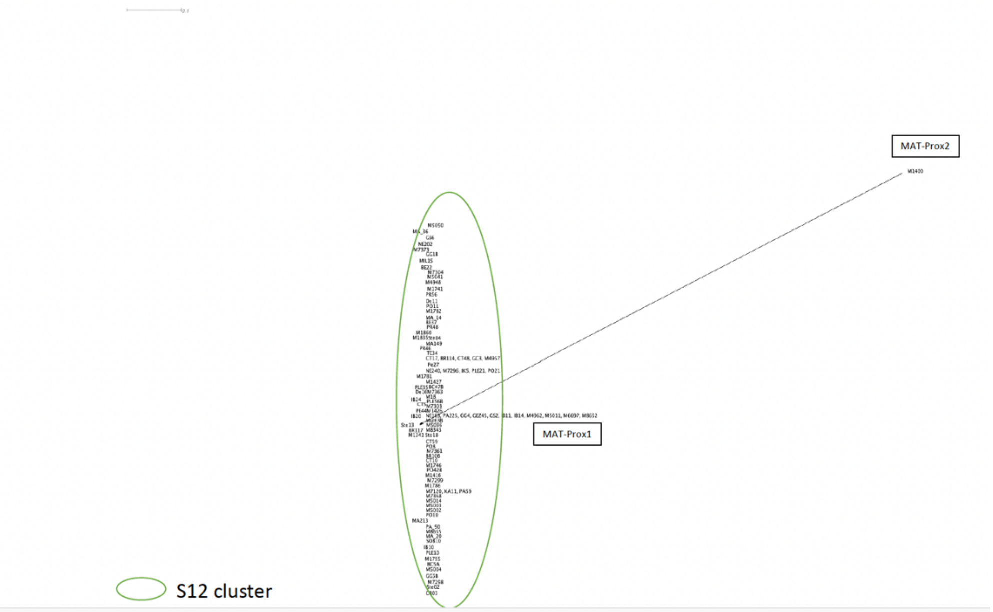


B (2) other contigs

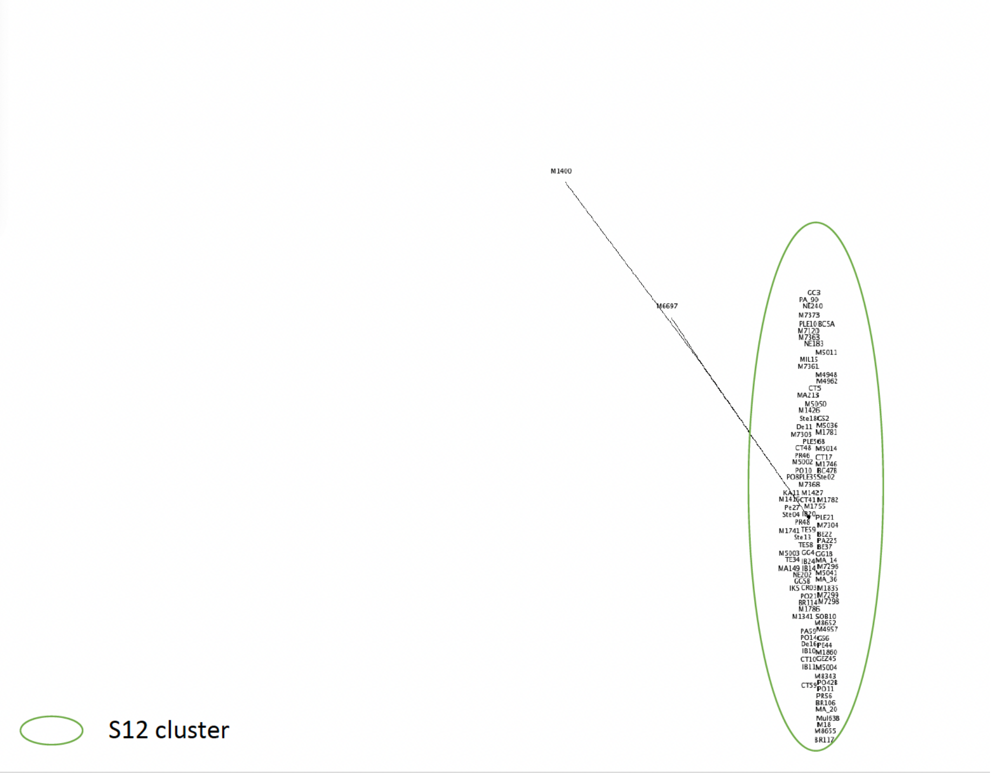

C (1) MAT-proximal region

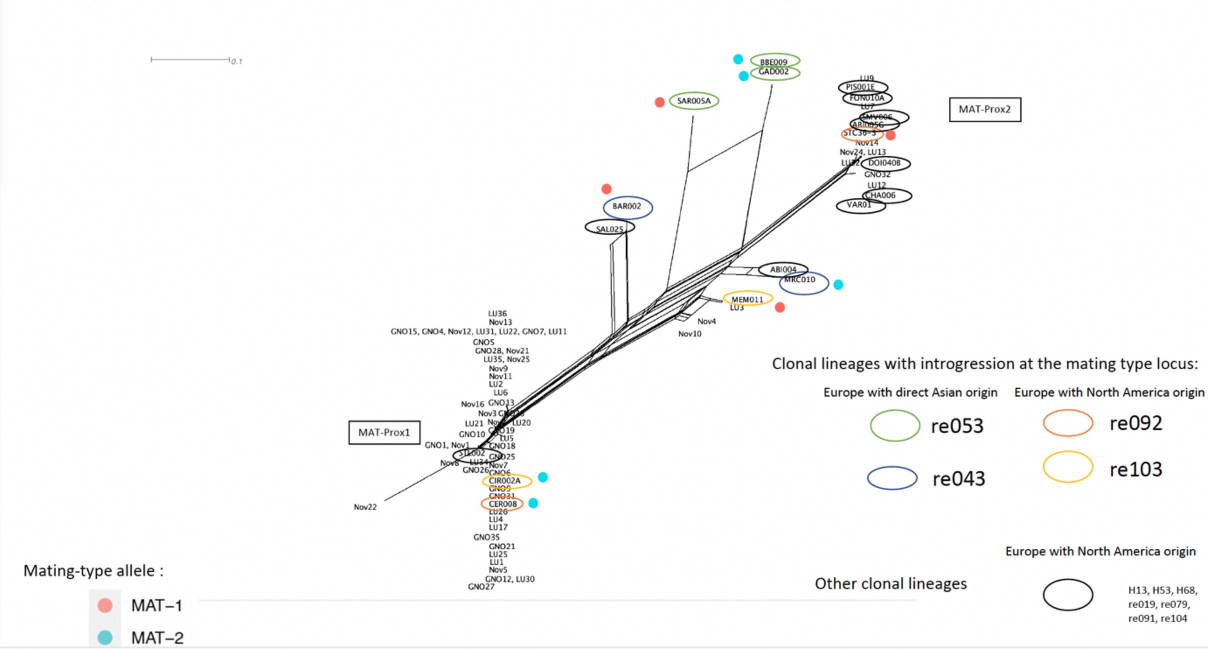


C (2) other contigs

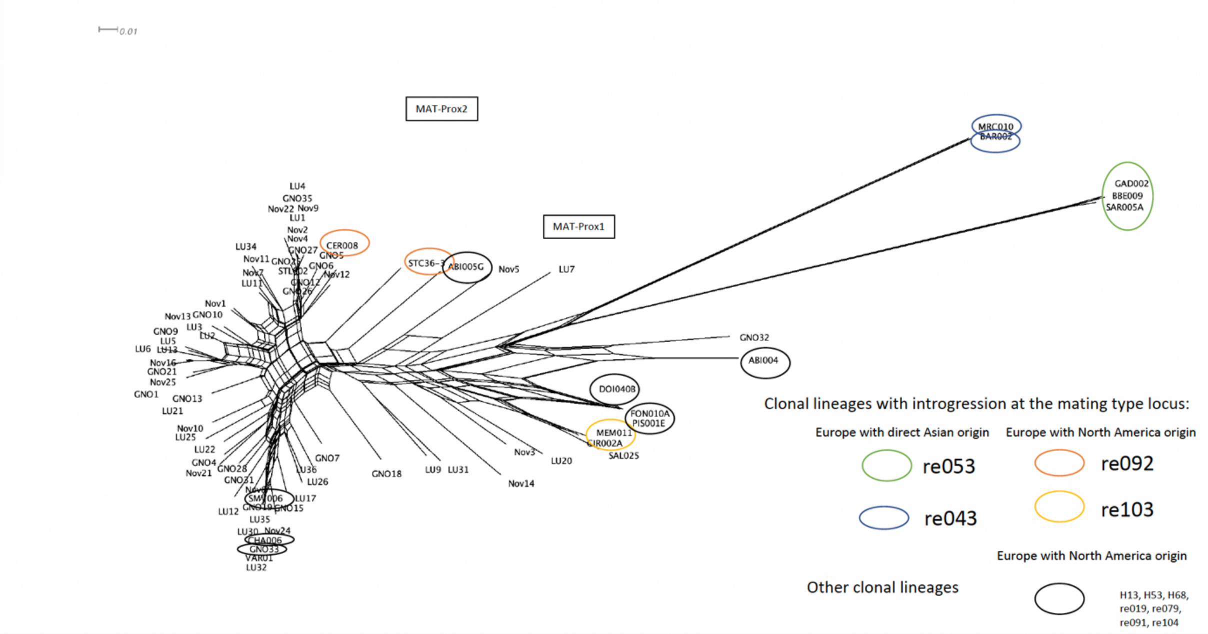

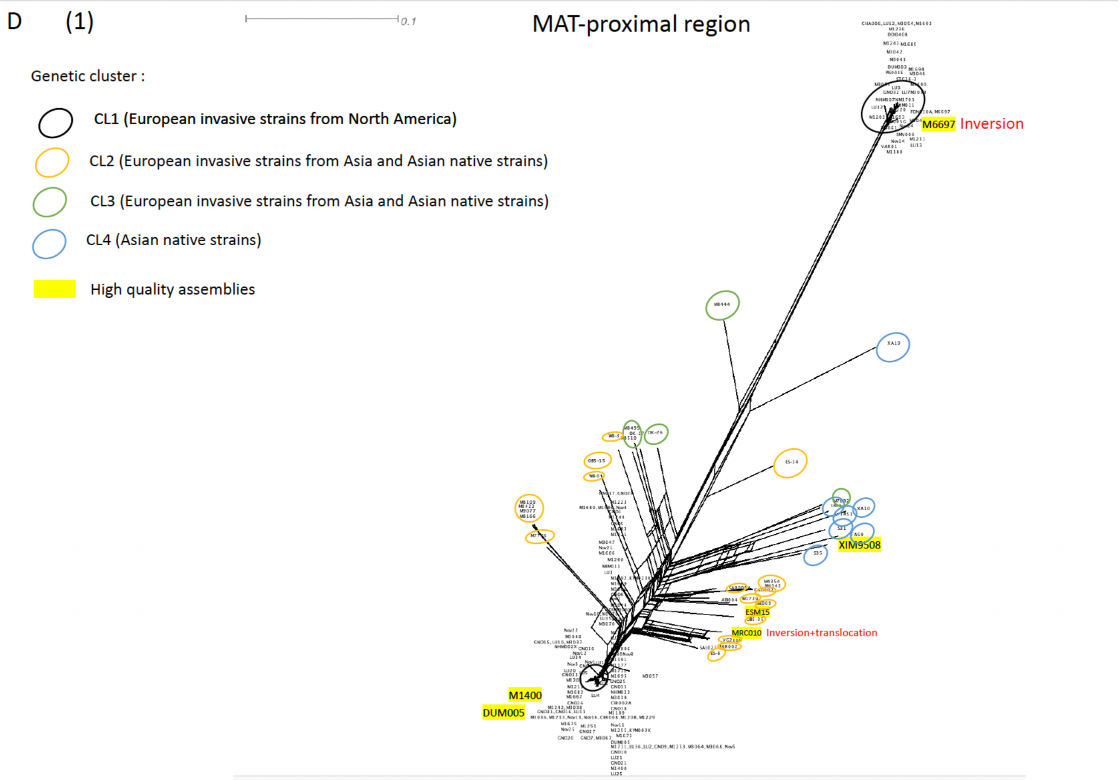
D (1)
0.1
MAT-proximal region
Genetic cluster :
CL1 (European invasive strains from North America)
CL2 (European invasive strains from Asia and Asian native strains)
CL3 (European invasive strains from Asia and Asian native strains)
CL4 (Asian native strains)
High quality assemblies
M6697 Inversion
XIM9508
ESM15
MRC010 Inversion+translocation
M1400
DUM005

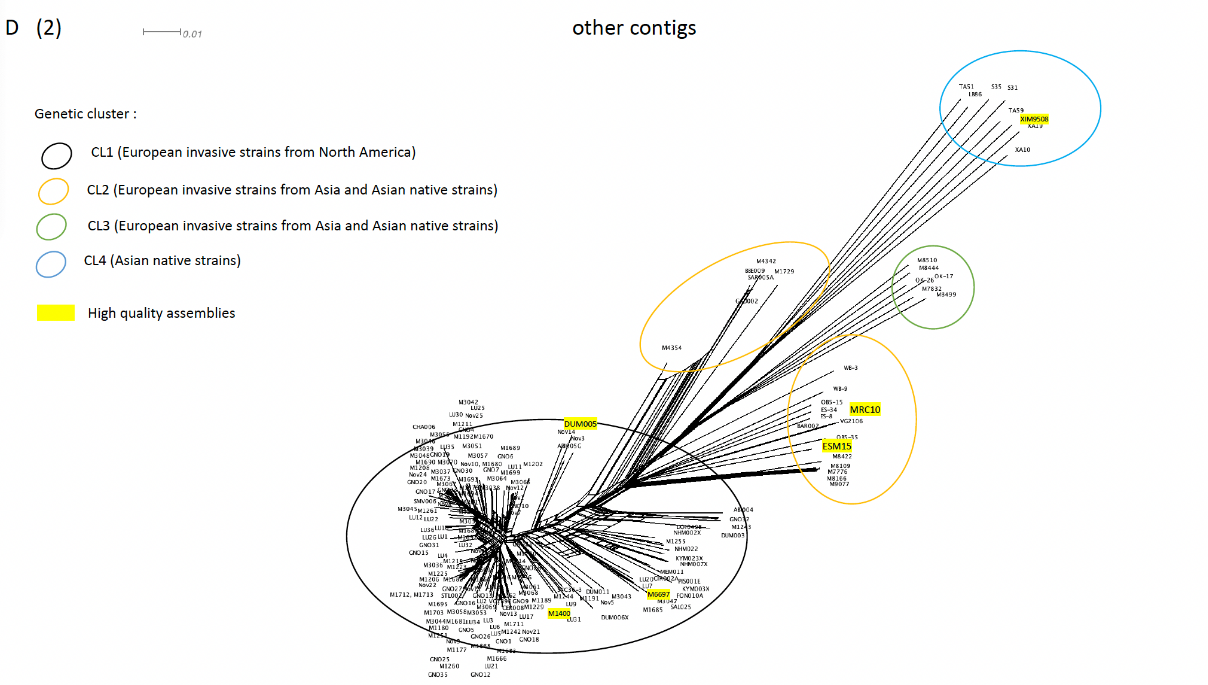
D (2)
0.01
other contigs
Genetic cluster :
CL1 (European invasive strains from North America)
CL2 (European invasive strains from Asia and Asian native strains)
CL3 (European invasive strains from Asia and Asian native strains)
CL4 (Asian native strains)
High quality assemblies
XIM9508
MRC10
ESM15
DUM005
M6697
M1400
